# ER-associated control of axonal vesicle trafficking during neuronal development

**DOI:** 10.64898/2026.04.23.720437

**Authors:** Mélody Subra, Janet B Meehl, Robert G Abrisch, Gia K Voeltz

## Abstract

Axonal vesicle trafficking is essential for presynaptic assembly, yet the intracellular structures that spatially and temporally regulate vesicle transport during neuronal development remain unclear. Here we identify previously uncharacterized membrane contact sites (MCSs) between the endoplasmic reticulum (ER) and both synaptic vesicle precursors (SVPs) and dense-core vesicles (DCVs). Using live confocal and cryo-electron microscopy, we reveal that vesicle transport undergoes a transient developmental slowdown in the axons of rat hippocampal neurons, associated with increased tripartite ER-vesicle-microtubule interfaces. Using proximity biotinylation, we identify the ER protein PRG1 as a developmentally regulated factor that modulates vesicle retention at these sites by interacting with KIF5A. Notably, a disease-associated PRG1 mutant fails to regulate vesicle trafficking. Finally, we demonstrate that PRG1 modulation of vesicle trafficking serves as a developmental checkpoint by delaying synapse formation and preventing network hyperactivity. Together, our findings establish ER-vesicle contact sites as a previously unrecognized layer of control of axonal trafficking and link intracellular organization to the timing of synapse assembly.

## Introduction

Neuronal communication depends on the precise assembly of functional synapses during development. A key step in this process is the timely delivery and accumulation of presynaptic components to nascent synaptic sites (Gundelfinger et al., 2016; Südhof, 2018; Zhai et al., 2001). In immature neurons, synaptic vesicle precursors (SVPs) and dense-core vesicles (DCVs) are continuously transported along axons to supply developing presynaptic terminals with neurotransmitters and neuropeptides required for future synaptic transmission (Dean et al., 2012; Guedes-Dias & Holzbaur, 2019). The regulation of axonal vesicle trafficking is therefore essential for synapse maturation and for the emergence of functional neuronal circuits (Rizalar et al., 2021). It remains unclear how vesicle mobility and accumulation are coordinated with the transition from a rapidly growing axon to a functionally mature presynaptic terminal. Understanding this regulation is critical, as premature or excessive vesicle delivery could alter synapse formation and is increasingly recognized as a pathogenic mechanism contributing to neurodevelopmental and neurodegenerative disorders (Sleigh et al., 2019; Xiong & Sheng, 2024).

Axonal vesicle transport is a dynamic process whereby SVPs and DCVs are generated in the soma and transported toward developing synaptic compartments. They undergo long-range bidirectional movements driven by kinesin and dynein motors along microtubules (MTs), interspersed with pauses and directional switches (Foster et al., 2022; Hirokawa & Takemura, 2005). These transport dynamics help determine how vesicles are distributed along axons and ultimately accumulate at presynaptic sites (Y. E. Wu et al., 2013). While the molecular machinery underlying vesicle mobility has been extensively characterized, much less is known about how vesicle trafficking is spatially and developmentally tuned to ensure proper timing of synapse assembly. One possibility is that axonal vesicle dynamics are regulated by ER membrane contact sites (MCSs), drawing parallels to other cytoplasmic organelles (Bonifacino & Neefjes, 2017; Jongsma et al., 2024; Mesmin et al., 2017; Raiborg, Wenzel, & Stenmark, 2015; Striepen & Voeltz, 2023). During early development, the ER adopts a distinctive ladder-like architecture that is required for axon growth and branching (Zamponi et al., 2022). Axonal ER ladders are composed of dynamic tubular rails that run the length of the axon and are interconnected by regularly spaced tubular rungs that wrap tightly around the MT bundle. Importantly, disruption of ER ladder integrity impairs SVP and DCV trafficking (Zamponi et al., 2022). While MCSs with other organelles have been visualized in neurons (Benedetti et al., 2025; Y. Wu et al., 2017), whether the ER directly interacts with vesicles involved in neurotransmission has not been investigated.

Here, we use live-cell imaging, reversible dimerization probes, and cryo-electron microscopy to visualize and describe ER-SVP and ER-DCV MCSs at early developmental stages in rat hippocampal neurons. We use proximity proteomics to uncover Plasticity-related gene 1 (PRG1) as an ER-localized protein enriched within these nano-environments. By combining developmental analyses with gain- and loss-of-function approaches, we reveal that PRG1 modulates axonal trafficking of SVPs, DCVs and late endosomes (LEs), through its association with the kinesin-1 motor KIF5A, and that PRG1 depletion enhanced early synapse assembly and increased neuronal activity. Together, these studies reveal a compelling cellular function for a clinically relevant protein, PRG1, that has previously been linked to epilepsy and neuronal hyperexcitability in human patients and mouse models (Knierim et al., 2023; Trimbuch et al., 2009; Vogt et al., 2016).

## Results

### SVPs and DCVs are transiently associated with the axonal ER ladder

The ER adopts a repetitive ladder like morphology during an early stage of neuronal development in growing axons (Zamponi et al., 2022). Disruption of axonal ER ladder morphology attenuates axon growth and the trafficking of SVPs and DCVs (Zamponi et al., 2022). We therefore asked whether the axonal ER interacts with trafficking vesicles to directly modulate their mobility during early stages of neuronal development. To determine whether contact sites exist between the axonal ER and SVPs and/or DCVs, we co-transfected primary rat hippocampal neurons at 4 days in vitro (DIV4) with a widely distributed luminal ER marker (mNeon-KDEL, green) and a synaptic vesicle (SV) resident protein (Synaptotagmin 1, Syt1-Halo, magenta) (Bradberry et al., 2020; Watson et al., 2023) or a well-established DCV protein (Synaptotagmin 4, Syt4-mNeon, magenta) (Bharat et al., 2017) used as vesicle markers. Neurons were visualized at DIV5 over one-minute live imaging sequences (Fig. 1a-f and movie S1). ER and SVP trajectories were compared using kymographs (Fig. 1c). While the ER ladder appeared largely stable over the time course of our videos, the SVPs moved bidirectionally along axons. Importantly, the SVPs frequently paused at ER rungs (orange arrows) consistent with previous reports (Zamponi et al., 2022). We also performed experiments to visualize the trajectories of DCVs relative to the ER. Time-lapse images and kymographs showed that DCVs also pause at ER rungs (Fig. 1d-f, orange arrows). In several recordings, ER and vesicle movements were coordinated, suggesting physical coupling between the two organelles (see kymographs, Fig. 1c,f). We measured the frequency of spatial overlap between vesicles and the ER (Fig. 1g,h). Approximately 80% of SVPs or DCVs colocalized with ER tubules, indicating a strong spatial correlation. As a control, we flipped the ER image horizontally to simulate random overlap; this significantly decreased co-localization suggesting that the association was not due to a random distribution (Fig. 1g,h).

**Figure 1.**
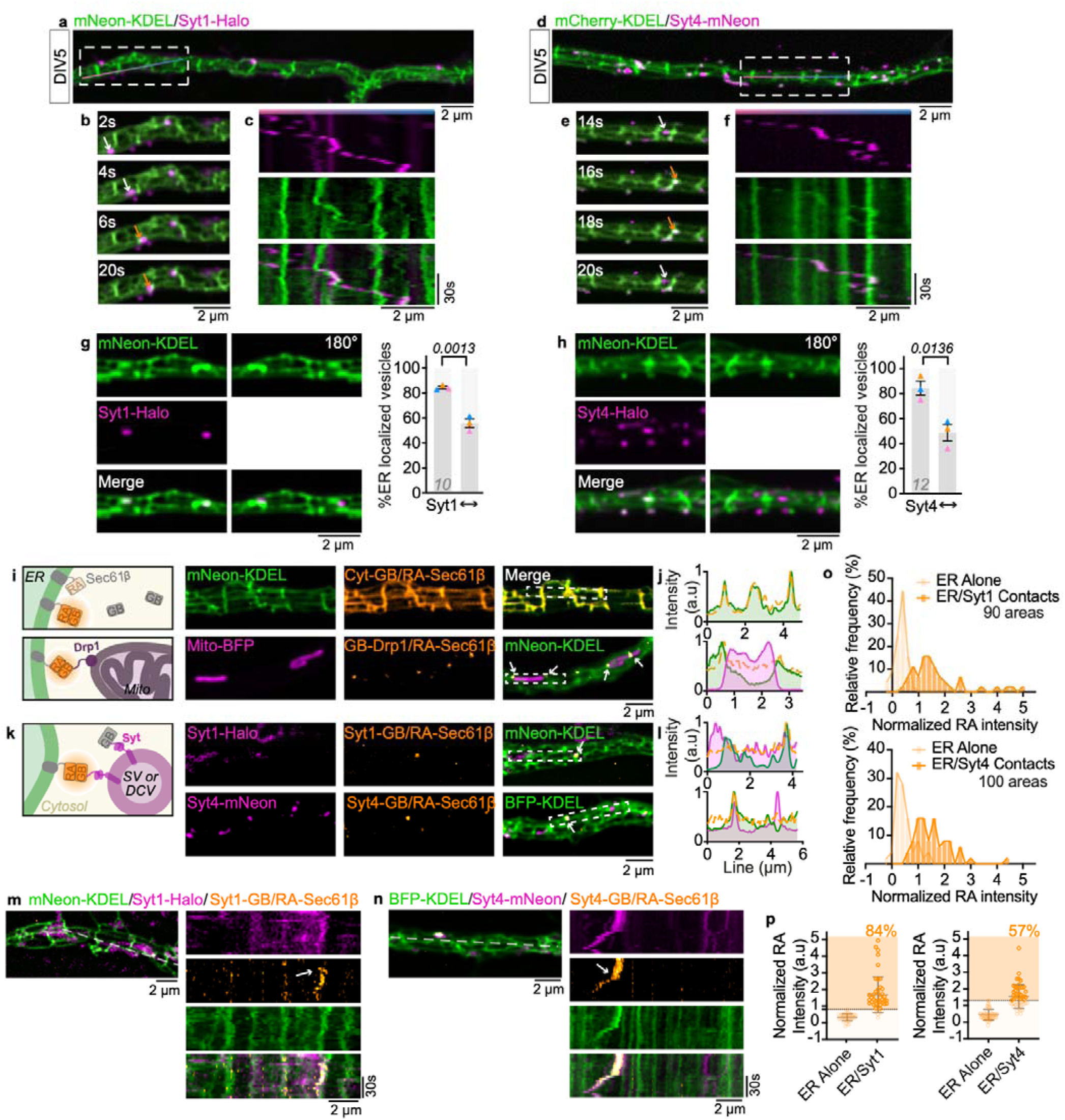
SVPs and DCVs are transiently associated with the axonal ER ladder. (a) Representative merged image of a DIV5 rat hippocampal neuron transfected with an ER luminal marker (mNeon-KDEL, green) and an SVP protein (Syt1-Halo, magenta). (b) Time-lapse corresponding to the rectangle drawn in (a) showing vesicle and ER dynamics over time (1 frame every 2.32 s). Orange arrows indicate vesicles closely associated with ER tubules; white arrows indicate moving vesicles. (c) Kymograph corresponding to the line drawn in (a), illustrating bidirectional SVP trafficking and frequent pauses at ER ladder sites resulting in ER ladder perturbation. (d,e,f) As in (a-c) performed with dense-core vesicles (DCVs) labeled with Syt4-mNeonGreen (magenta), showing similar dynamic association with ER tubules (mCherry-KDEL, green). (g,h) Quantification of the fraction of vesicles (Syt1 and Syt4, respectively, in magenta) whose signal overlaps with the resolvable ER tubules (green), compared to a control condition in which the ER region of interest (ROI) was flipped horizontally, revealing background-level overlap. Colocalization was quantified across at least 30 ROIs from 10 to 12 cells per condition, n=3 independent experiments. Data shown are mean ± SEM; unpaired t-tests. (i) Schematic and representative images of the control ddFP system used in DIV5 neurons, with RA-Sec61β at the ER and a cytosolic GB or a mitochondrial protein GB-Drp1. Dimerization of RA and GB domains increases RA-fluorescent signal (orange). Cells were also cotransfected with an ER (green) and mitochondrial marker (magenta). The spatial overlap between the ddFP signal and the ER-mitochondria contact sites is indicated by white arrows. (j) Line-scan intensity profile across (i) images, illustrating the spatial overlap between the ddFP signal (GB-Drp1) relative to the ER-mitochondria contact sites. (k) Schematic and representative images of the ddFP system used to assay ER-vesicle contacts. ddFP fluorescence intensity is increased (in orange) at locations where RA-Sec61β (ER, green) and GB-tagged Syt1 (SVPs, magenta) or Syt4 (DCVs, magenta) are closely apposed (white arrows). (l) Line-scan intensity profile across the contact site shown in (k), illustrating spatial overlap of ER, vesicle, and ddFP signal. (m,n) Kymographs showing ddFP contact sites (orange) relative to locations (at white arrow) where (m) Syt1- or (n) Syt4-labeled vesicles encounter the ER (time course: 1 frame every 3.13 s). (o) Frequency distributions of ddFP intensity values taken from regions of ER alone (monomer) and regions where ER crossed a Syt1- or Syt4-vesicle (dimer) validate signal increase upon dimerization (n = 90 to 100 spots pooled from 2 biological replicates). (p) Normalized ddFP intensities at ER-only and ER-vesicle crossing regions. ER-vesicle crossings with ddFP intensity exceeding the maximum intensity measured in ER-only regions were classified as ddFP-positive contact sites (above dotted line), identifying 84% of Syt1 and 57% of Syt4 ER-vesicle crossings as bona fide MCSs.

We next used the dimerization-dependent fluorescent protein (ddFP) system (Ding et al., 2015) to visualize whether the ER and vesicles come within MCS tethering distances in living cells. The ddFP system consists of a green-binding (GB) domain and a red-acceptor (RA) domain. Dimerization of GB and RA domains increases red fluorescence, and for our purposes indicates proximity within the MCS size range (10-30nm) (Abrisch et al., 2020; Lee et al., 2020). A relevant feature of the ddFP system is that it is reversible (Kd ∼7 uM) (Ding et al., 2015), thus enabling dynamic visualization of organelle contacts in real time. To validate this approach for scoring MCSs in neurons, we co-transfected cells with Sec61β-RA and GB-fusions to either a cytosolic protein (Cyt-GB) or an ER-mitochondria MCS protein (GB-Drp1, Dynamin Related Protein 1) (Fig. 1i). These dimer pairs fluoresced on the ER and at ER-mitochondrial MCSs, respectively, as predicted (Fig. 1i,j, arrows). Next, we co-transfected neurons with Sec61β-RA and Syt1-GB or Syt4-GB to visualize contacts between the ER and SVPs or DCVs (Fig. 1k,l). Discrete positions of contact could be observed (see arrow marking yellow “ddFP” puncta in Fig. 1k). Quantification of ddFP fluorescence revealed a clear enrichment of signal at ER-vesicle interfaces compared with ER-only regions, with 84% of ER-associated SVPs and 57% of ER-associated DCVs displaying detectable ddFP signals (Fig. 1o,p). One-minute live recordings further revealed that ddFP-positive contacts could remain stationary or move in coordination with ER tubules, occasionally accompanying vesicle pausing or transport (Fig. 1m,n). Together, these observations reveal that SVPs and DCVs form stationary and dynamic contact sites with the axonal ER during early development and that these contacts might modulate their trafficking behavior.

### Proximity proteomics identifies PRG1 as a regulator of vesicular trafficking

We performed proximity-dependent biotinylation experiments in rat hippocampal neurons to identify the molecular components of ER contact sites with SVPs and DCVs. Our strategy was to fuse the TurboID biotin ligase to the C-terminal domain of Syt1-mNeon-V5, Syt4-mNeon-V5, and compare ER proteins biotinylated by these constructs to a cytosolic control (mNeon-V5-TurboID, empty vector, EV) (Fig. 2a). TurboID is well suited for identifying MCS proteins because it catalyzes the biotinylation of lysine residues within MCS distances (10–30 nm) (Branon et al., 2018; Hoyer et al., 2018; Nguyen & Voeltz, 2022; Roux et al., 2012). Western blot and immunofluorescence analysis confirmed efficient labeling: both Syt1-mNeon-V5-TurboID and Syt4-mNeon-V5-TurboID colocalized with biotinylated proteins detected using Streptavidin-AF555 (Fig. 2b and S1a,b). Mass spectrometry analysis identified numerous close neighbors of both Syt1 and Syt4 (a few examples are highlighted in orange, Fig. 2c,d), validating the specificity of the assay. Comparative analysis between the SVP and DCV datasets revealed plasticity-related gene 1 (PRG1) as a shared candidate enriched in both interactomes (Fig. 2c,d).

**Figure 2.**
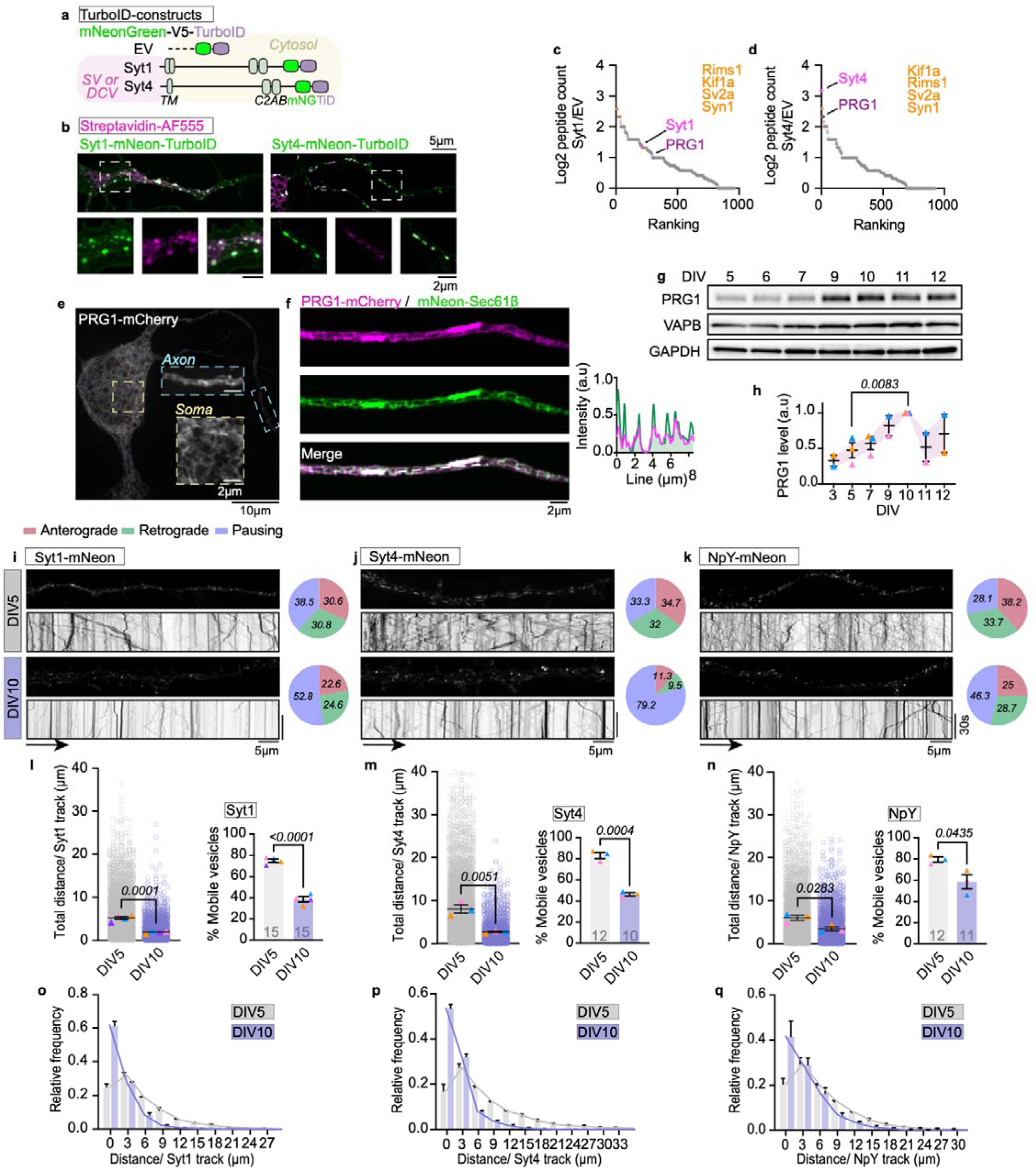
Proximity proteomics identifies PRG1 as a regulator of vesicular trafficking. (a) Schematic of the TurboID proximity labeling strategy. Syt1-mNeon-V5-TurboID, Syt4-mNeon-V5-TurboID or an empty-vector (EV) control (mNeon-V5-TurboID) construct were expressed in rat hippocampal neurons to identify ER proteins near SVPs and DCVs, respectively. Neurons were incubated with biotin for 15 min, and biotinylated proteins were purified and analyzed by mass spectrometry (MS). (b) Representative images showing biotinylated proteins detected by streptavidin-AF555 staining in neurons expressing either Syt1-mNeon-V5-TurboID (left panel) or Syt4-mNeon-V5-TurboID (right panel) following a 15-min biotin incubation. The streptavidin signal overlaps with the vesicle markers, confirming specificity. (c,d) Rank plots showing proteins biotinylated by Syt1-mNeon-V5-TurboID or Syt4-mNeon-V5-TurboID relative to EV controls. Hits include Syt1 and Syt4 (in magenta), previously published proximal proteins (in orange), and PRG1 (in purple). (e) Representative image of PRG1-mCherry distribution in the soma and the axonal segment (ROI indicated) of a DIV5 hippocampal neuron. (f) Representative images and corresponding line-scan intensity profiles confirm colocalization of PRG1-mCherry with a general ER marker (mNeon-Sec61β) in DIV5 hippocampal neurons. (g) Immunoblot analysis of endogenous protein levels for PRG1, VAPB, and GAPDH across rat hippocampal neuron development (DIV5-DIV12). (h) Quantification of PRG1 protein levels in (g) normalized to the maximum from 3 independent cultures. Data are mean ± SEM; unpaired t-test. (i,j,k) Representative images and corresponding kymographs of (i) Syt1-mNeon, (j) Syt4-mNeon or (k) NpY-mNeon vesicle dynamics in DIV5 (top panels) or DIV10 (bottom panels) hippocampal neurons. Each line corresponds to the track of an individual vesicle. Pie charts indicate the proportion of anterograde, retrograde and pausing events (see Methods and Fig S1j). Values indicated in the pie charts represent the mean proportion of events across all neurons for which axonal orientation could be determined, from 3 to 4 independent experiments. Note that SVP and DCV mobility is reduced at DIV10 compared with DIV5. (l,m,n) Quantification of total distance traveled and associated mobile fraction of tracks of (l) Syt1-mNeon, (m) Syt4-mNeon and (n) NpY-mNeon labeled-vesicles at DIV5 versus DIV10. Vesicles were classified as mobile or immobile using a displacement threshold of 1.5 µm. Data are mean ± SEM from at least n=3 independent experiments, 11 to 15 neurons were analyzed per condition. The precise number of cells analyzed is indicated on each graph. Statistics: unpaired t-test. (o,p,q) Relative frequency distributions of the distance traveled by individual (o) Syt1-mNeon, (p) Syt4-mNeon and (q) NpY-mNeon vesicles at DIV5 and DIV10. Data are pooled from the neurons quantified in (l–n).

PRG1 belongs to the lipid phosphate phosphatase (LPP)–related protein family, although it lacks confirmed phosphatase activity (Bräuer et al., 2003; McDermott et al., 2004; Tang et al., 2015). Earlier studies described PRG1 at the plasma membrane, where it regulates lysophosphatidic acid (LPA) signaling at glutamatergic synapses and PPP2A/β1-integrin signaling influencing neuron survival, growth and plasticity (Hashimoto et al., 2013; X. Liu et al., 2016; Polyzou et al., 2024; Vogt et al., 2016). However, our live-cell imaging experiments revealed that wild-type PRG1-mCherry co-localized well with an ER protein (mNeon-Sec61β) in the soma and axonal compartments of rat hippocampal neurons (Fig. 2e,f). Fluorescently tagged PRG1 constructs also co-localize with ER markers in tissue culture cells (see COS-7 and HEK293T, Fig. S1c,d) irrespective of tag identity (mCherry or mNeonGreen) or location (N or C terminus). According to several algorithms, PRG1 is a type II membrane protein featuring an N-terminal six-pass transmembrane domain and a ∼400 amino-acid cytosolic unstructured C-terminal tail (Fuchs et al., 2022). The cytosolic tail contains a conserved, high-affinity, Ca²⁺-dependent calmodulin-binding site for which the function is unknown (Tokumitsu et al., 2010)(Fig. S1c).

We analyzed the temporal expression of endogenous PRG1 in embryonic rat hippocampal neurons cultured from 4 to 14 days in vitro (DIV4-DIV14). Immunoblot and mRNA analyses revealed that PRG1 levels are low at early stages, increase progressively until DIV10, and then slightly decline before reaching a plateau (Fig. 2g,h and Fig. S1e,f). This developmental profile is consistent with previous reports describing PRG1 mRNA upregulation during hippocampal maturation (Gross et al., 2022), as well as with expression patterns from rodent and human brain atlases (Cardoso-Moreira et al., 2019)(Fig. S1h,i). When compared to reference developmental markers such as Synaptophysin (Syp) and Neurog2, the DIV5–DIV10 window in rats roughly corresponds to the perinatal stage in humans (Fig. S1h,i).

If PRG1 is regulating vesicle trafficking, SVP and DCV mobility should differ between developmental stages. To test this hypothesis, primary rat hippocampal neurons were transfected with Syt1-mNeon or Syt4-mNeon 24 hours prior to imaging and 45 second live recordings were collected at successive time points. Vesicle mobility was quantified as the total distance traveled by each vesicle (*TrackMate*, Fiji) (Fig. 2i-q and movie S2). SVP and DCV trajectories were also compared using kymographs at DIV5 and DIV10 (Fig. 2i,j). Individual trafficking events were classified as anterograde, retrograde, or pausing based on their directionality and displacement (Fig. S1j). At DIV5, vesicular transport was highly dynamic, characterized by bidirectional long and continuous runs. By DIV10, vesicle movement was markedly reduced, with fewer mobile events (Fig. 2i,j). This reduction was not associated with a directional bias toward anterograde or retrograde transport. The distribution of individual track distances also shifted toward shorter trajectories at DIV10 (Fig. 2o,p), before partially recovering at later stages (Fig. S1g). *Trackmate* analysis confirmed that the mobile fraction of SVPs and DCVs decreased from approximately 75% to 40% and 80% to 50%, respectively (Fig. 2l,m) with a notable increase in vesicle clustering at DIV10 (Movie S2). Similar results were obtained when Neuropeptide-Y (NpY), a canonical cargo of DCVs, was used as an additional marker in these experiments (Fig. 2k,n,q) (Persoon et al., 2018). Together, our data provide the first description of a transitional reduction in vesicular trafficking during a period of hippocampal neuronal development. Notably, this period correlates with higher levels of PRG1.

### An epilepsy-associated PRG1 mutation abolishes vesicle trafficking control

We hypothesized that ER-localized PRG1 might alter vesicle mobility during the early stages of neuronal development. We therefore tested if overexpression of PRG1 at DIV5, a developmental stage when vesicle trafficking is high and PRG1 levels are low, would be sufficient to reduce vesicle mobility. Hippocampal neurons were co-transfected at DIV4 with either PRG1-mCherry or mCherry-Sec61β (control) and with Syt1-mNeon (SVPs), Syt4-mNeon or NpY-mNeon (DCVs) to visualize the effect of PRG1 expression on SVP and DCV mobility, respectively (Fig. 3 and S2). Vesicular trafficking was quantified at DIV5 by acquiring live-cell movies and tracking single vesicles under each condition (*TrackMate*, Fiji). These movies revealed that PRG1 overexpression markedly reduced vesicle trafficking relative to Sec61β-expressing controls (Fig. 3c-f, S2a and movie S3). Kymographs demonstrated that both anterograde and retrograde transport were being affected (Fig. 3c,d). Further quantifications confirmed a significant decrease in run length, with a reduction of the mobile fraction from roughly 80% to 20% for SVPs and 85% to 40% for DCVs (Fig. 3e,f and S2b,c). Thus, PRG1 overexpression is sufficient to reduce vesicle mobility at DIV5.

**Figure 3.**
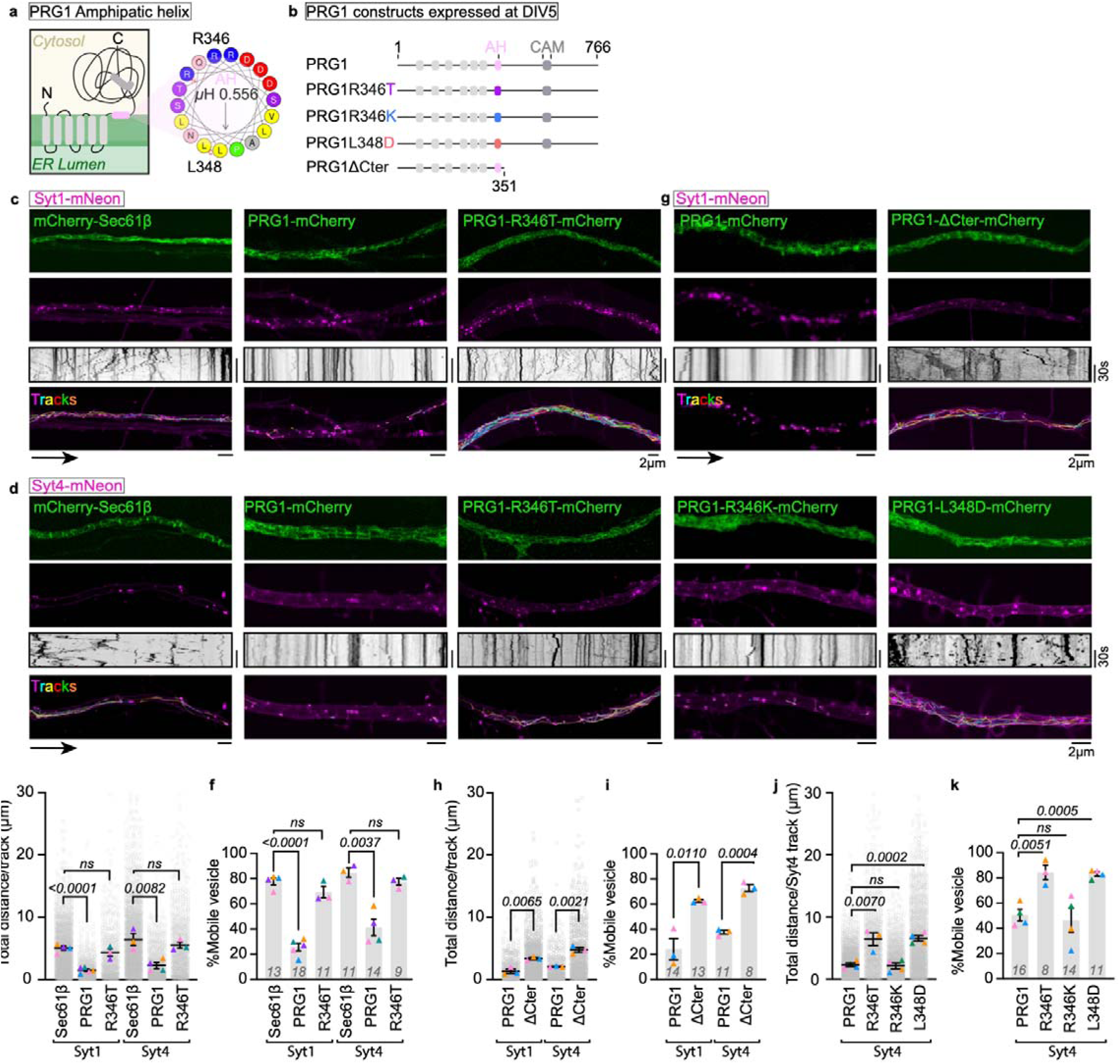
PRG1 overexpression inhibits vesicular trafficking; the epilepsy-associated mutant R346T is functionally inactive. (a) Schematic representation of PRG1 domain organization. The amphipathic helix (AH, residues 342-359: RDALRSLTDLNQDPSRVL, in magenta) is adjacent to the last transmembrane domain. (b) PRG1 constructs tested, including mutations at positions R346 (R346T and R346K) and L348 (L348D), and a C-terminal deletion (ΔCter). (c) Representative images, kymographs and TrackMate-generated tracks from 45 s movies of Syt1-mNeon labeled vesicles (magenta) in DIV5 hippocampal neurons expressing either mCherry-Sec61β (ER), PRG1-mCherry, or PRG1-R346T-mCherry (in green). Note that PRG1-mCherry overexpression reduces vesicle mobility whereas the PRG1-R346T-mCherry epilepsy-associated mutant does not. (d) Representative images, kymographs and TrackMate-generated tracks from recordings of Syt4-mNeon labeled vesicles (magenta) in DIV5 hippocampal neurons expressing either mCherry-Sec61β (ER), PRG1-mCherry, or PRG1-R346T-mCherry (in green). Syt4 vesicle trafficking was additionally assessed in neurons expressing PRG1-R346K-mCherry or PRG1-L348D-mCherry. Compared with wild-type PRG1, the R346K mutant retains the ability to reduce vesicle trafficking, whereas the L348D mutant does not. (e,f) Quantification of (e) the total distance traveled and (f) mobile fraction for data from (c) and (d). Data are mean ± SEM from n=3 to 4 independent experiments compared through unpaired t-tests, 9 to 18 neurons were analyzed per condition. (g) Representative images, kymographs and TrackMate-generated tracks from recordings of Syt1-mNeon labeled vesicle dynamics in DIV5 neurons expressing PRG1-mCherry and PRG1-ΔCter-mCherry. Note that compared to wild-type PRG1, the mutant does not reduce vesicle trafficking. (h,i) Quantification of (h) total distance traveled and (i) mobile fraction of Syt1- or Syt4-mNeon-labeled vesicles in DIV5 neurons expressing PRG1-mCherry or PRG1-ΔCter-mCherry. Data are mean ± SEM from n=3 independent experiments compared using unpaired t-tests, 8 to 14 neurons were analyzed per condition. (j,k) Quantification of (j) total distance traveled and (k) mobile fraction of images in experiment (d). Data are mean ± SEM from at least n = 3 experiments compared through unpaired t-tests, 11 to 16 neurons were analyzed per condition. Eight additional cells expressing PRG1-R346T-mCherry alongside Syt4-mNeon, as in (e-f), were analyzed in this series of experiments.

PRG1 has previously been linked to epilepsy and neuronal hyperexcitability in human patients and mouse models (Knierim et al., 2023; Trimbuch et al., 2009). A pathogenic missense mutation (R346T) located within the first α-helix of the C-terminal domain of PRG1 has been identified in juvenile epileptic patients and replicated in mouse models (Knierim et al., 2023; Trimbuch et al., 2009; Vogt et al., 2016) (Fig. 3a). This mutation constitutes a loss-of-function variant that is unable to rescue the neuronal hyperexcitability observed in PRG1 knockout neurons, although the molecular mechanism remains unresolved. The R346 residue exists within a highly conserved α-helix proposed by different structural predictors to be positioned in the cytosol adjacent to PRG1’s last transmembrane domain (Fig. 3a,b). To gain insight into the mechanism of this disease mutant at the molecular level, we tested whether overexpression of the epilepsy-associated PRG1 mutant (PRG1-R346T-mCherry) could reduce vesicle mobility. The PRG1-R346T mutant localized to the ER similarly to wild-type PRG1 (Fig. 3c,d) but failed to alter vesicular transport, showing no significant change in displacement or mobility compared with mCherry-Sec61β expressing neurons (Fig. 3c-f and S2a-c).

HeliQuest analysis predicted the helix carrying the pathological mutation to be amphipathic (AH) (Gautier et al., 2008)(Fig. 3a). We hypothesized that this AH could contribute to PRG1-dependent vesicle trafficking regulation. To test this idea, we generated two additional point mutants. An R346K substitution alters the same residue as the pathological R346T variant but preserves the amphipathic character of the helix. This mutant reduced vesicular trafficking similar to wild type PRG1. In contrast, an L348D mutation that should abolish the hydrophobic face of the helix and disrupt its amphipathic nature (Fig. 3a,b) did not alter vesicle dynamics (Fig. 3d,j,k). To further investigate the contribution of the cytosolic region, we generated a C-terminus truncation mutant (PRG1-ΔCter, residues 1-351) that retains the amphipathic helix but lacks the downstream cytosolic domain (Fig. 3b). Despite its ER localization, PRG1-ΔCter-mCherry failed to reduce SVP or DCV mobility (Fig. 3g-i), indicating that the amphipathic helix alone is not sufficient to regulate trafficking. Importantly, PRG1-mCherry and its mutants showed comparable expression levels without inducing obvious alteration in overall neuronal or ER morphology (Fig. S2d-h). We therefore propose that both the amphipathic character of this cytosolic α-helix and the downstream C-terminal region are required for PRG1-dependent trafficking regulation.

### PRG1 depletion rescues SVP and DCV trafficking at DIV10

Having established that PRG1 overexpression is sufficient to reduce vesicle mobility at DIV5, we next examined whether endogenous PRG1 contributes to the developmental trafficking change observed at DIV10. Neurons were infected with a PRG1-targeting shRNA (shPRG1-mCherry) at DIV4 and were transfected with SVP and DCV expression constructs at DIV8 and analyzed at DIV10. Western blot and qPCR analyses confirmed reduction of PRG1 expression under knockdown (KD) conditions compared to infection with a control short hairpin RNA (shCT-mCherry) (Fig. 4a-c). At DIV10, only mCherry-positive neurons were analyzed to restrict the analysis to PRG1-depleted neurons. Live-cell imaging of vesicles labeled with Syt1-mNeon (Fig. 4d), Syt4-mNeon (Fig. 4e), and NpY-mNeon (Fig. 4f) in DIV10 neurons treated with shPRG1-KD were significantly more mobile than shCT control cells. Kymographs also revealed longer vesicle trajectories following PRG1 depletion (Fig. 4d-f and movie S4). Quantitative analysis indicated significant increases in the total distance traveled per vesicle and the mobile fraction for both SVPs and DCVs in PRG1-KD neurons at DIV10 (Fig. 4g-i, m-o). Accordingly, the distributions of individual track distances shifted toward longer trajectories following PRG1 depletion (Fig. 4j-l). Directionality analysis further revealed a reduction in pausing events, with increased proportions of both anterograde and retrograde movements (Fig. 4d-f and m-o), mirroring the trafficking pattern observed at DIV5 (Fig. 2i-k). Interestingly, the maximum velocity of the remaining motile vesicles was not significantly altered by PRG1 depletion, suggesting that PRG1 primarily regulates vesicle pausing rather than the intrinsic speed of motor-driven transport (Fig. S3a). Thus, PRG1 depletion reverses the developmental alteration of vesicle trafficking observed around DIV10 in rat hippocampal neurons, coinciding with the developmental peak of endogenous PRG1 expression. Together, these data establish PRG1 as a regulator of axonal trafficking, promoting vesicle pausing during a specific developmental period.

**Figure 4.**
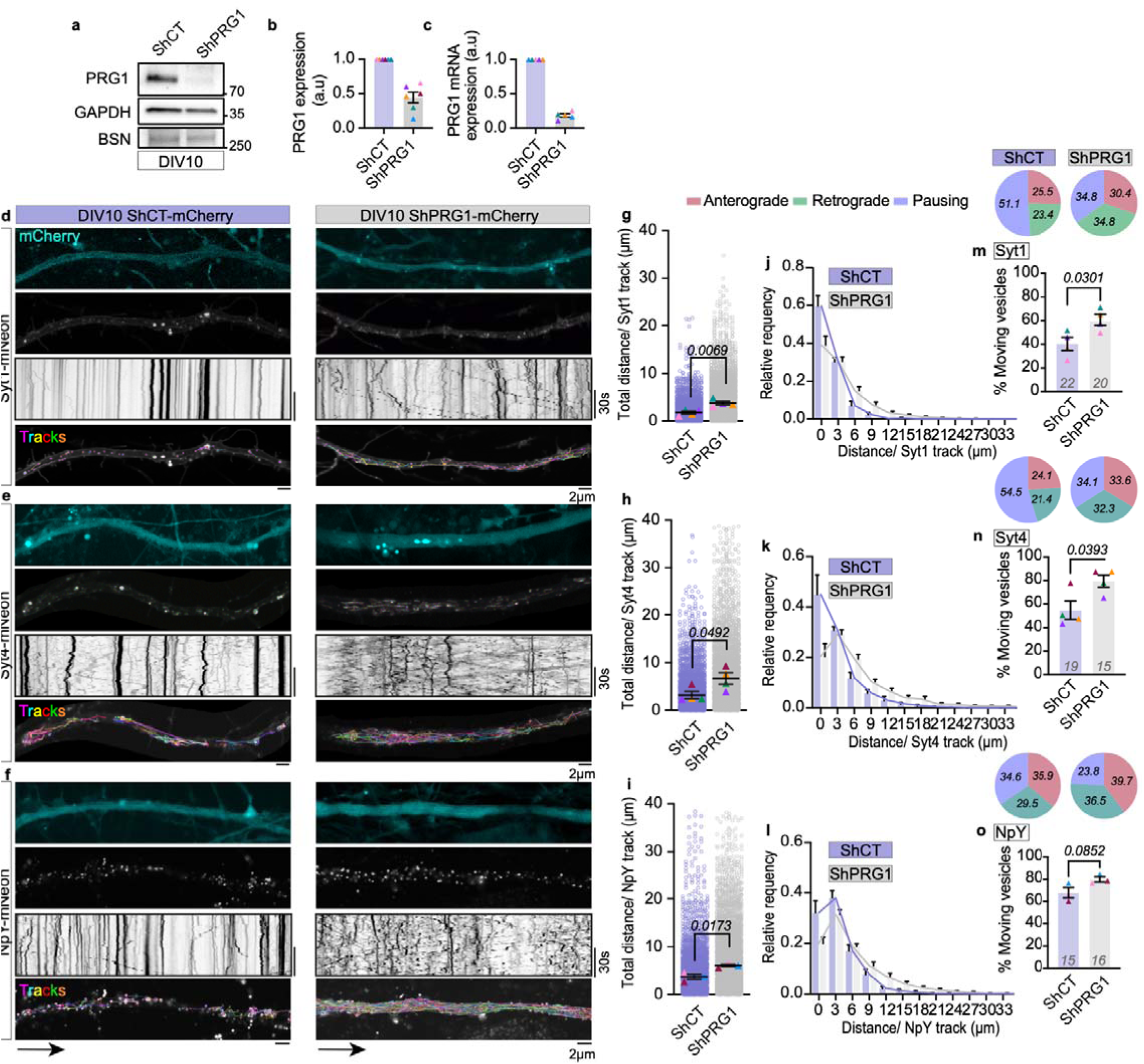
PRG1 depletion rescues SVP and DCV trafficking at DIV10. (a) Immunoblot analysis shows reduced PRG1 protein levels in shPRG1-infected neurons compared to shCT controls. Bassoon (BSN) and GAPDH are unchanged. (b,c) Quantitative analyses of immunoblot and qPCR data confirm that PRG1 protein and mRNA levels are reduced in shPRG1-treated neurons compared to controls. Data are mean ± SEM from 6 independent knockdown experiments used in Figure 4 and 7. (d,e,f) Representative images, kymographs and TrackMate-generated tracks from DIV10 hippocampal neurons expressing (d) Syt1-mNeon, (e) Syt4-mNeon and (f) NpY-mNeon (gray) under control or shPRG1 knockdown conditions, showing increased vesicle mobility upon PRG1 depletion. ShRNA-mCherry cassette is shown in cyan. (g,h,i) Quantification of experiments (d-f) shows a significant increase in the total distance traveled by vesicles in PRG1-depleted neurons. Data are mean ± SEM from at least n=3 independent experiments compared using unpaired t-tests. 20 to 22 neurons were analyzed per condition for Syt1-mNeon tracks and 15 to 19 neurons for Syt4- or NpY-mNeon tracks. (j,k,l) Relative frequency distributions of the distance traveled by individual (j) Syt1-mNeon-, (k) Syt4-mNeon- and (l) NpY-mNeon-labeled vesicles in control and PRG1-depleted neurons. Data are pooled from the neurons quantified in (g-i). (m,n,o) Quantification of the mobile fraction for (m) Syt1-mNeon-, (n) Syt4-mNeon- and (o) NpY-mNeon-labeled vesicles in control and PRG1-depleted neurons. Data are mean ± SEM from at least 3 independent experiments. Mobile fractions were compared using unpaired t-tests. Pie charts show the proportion of anterograde, retrograde and pausing events (see Methods and Fig S1j). Values indicated in the pie charts represent the mean proportion of events across all neurons for which axonal orientation could be determined, from 3 to 4 independent experiments.

### PRG1 complexes with KIF5A and Protrudin to pause late endosome trafficking

To investigate how PRG1 regulates axonal vesicle trafficking, we immunoprecipitated endogenous PRG1 from DIV10 hippocampal neurons to identify associated proteins. Mass spectrometry identified a large number of peptides from the kinesin-1 family (KIF5A/B/C) an interaction that was also confirmed for Kif5A by immunoblot analysis (Fig. 5a-c). We turned to HEK293T cells to discriminate whether PRG1 binds one of the KIF5 paralogs preferentially. PRG1-mCherry was overexpressed with a myc-tagged version of each of the KIF5 paralogs (KIF5A-C). HEK293T extracts were collected and PRG1-mCherry and associated proteins were purified on mCherry-trap beads. Immunoblot analysis was used to compare the bound KIF5 paralogs. These data revealed a stronger association of PRG1 with KIF5A than with KIF5B or KIF5C (Fig. 5d,e). To map the interacting region, we generated a series of KIF5A truncation constructs. PRG1 co-purified more of the KIF5A mutant lacking its motor domain (339-1032) than the full-length KIF5A and deletion of the C-terminal tail reduced this interaction (Fig. 5f,g). These data indicate that PRG1 preferentially associates with a KIF5A that lacks the motor domain, favoring its open conformation (Carrington et al., 2024; Kaan et al., 2011). These data are reminiscent of the ER contact site protein Protrudin that binds similarly to KIF5A (Matsuzaki et al., 2011) to promote late endosome (LE) trafficking (Raiborg, Wenzel, Pedersen, et al., 2015).

**Figure 5.**
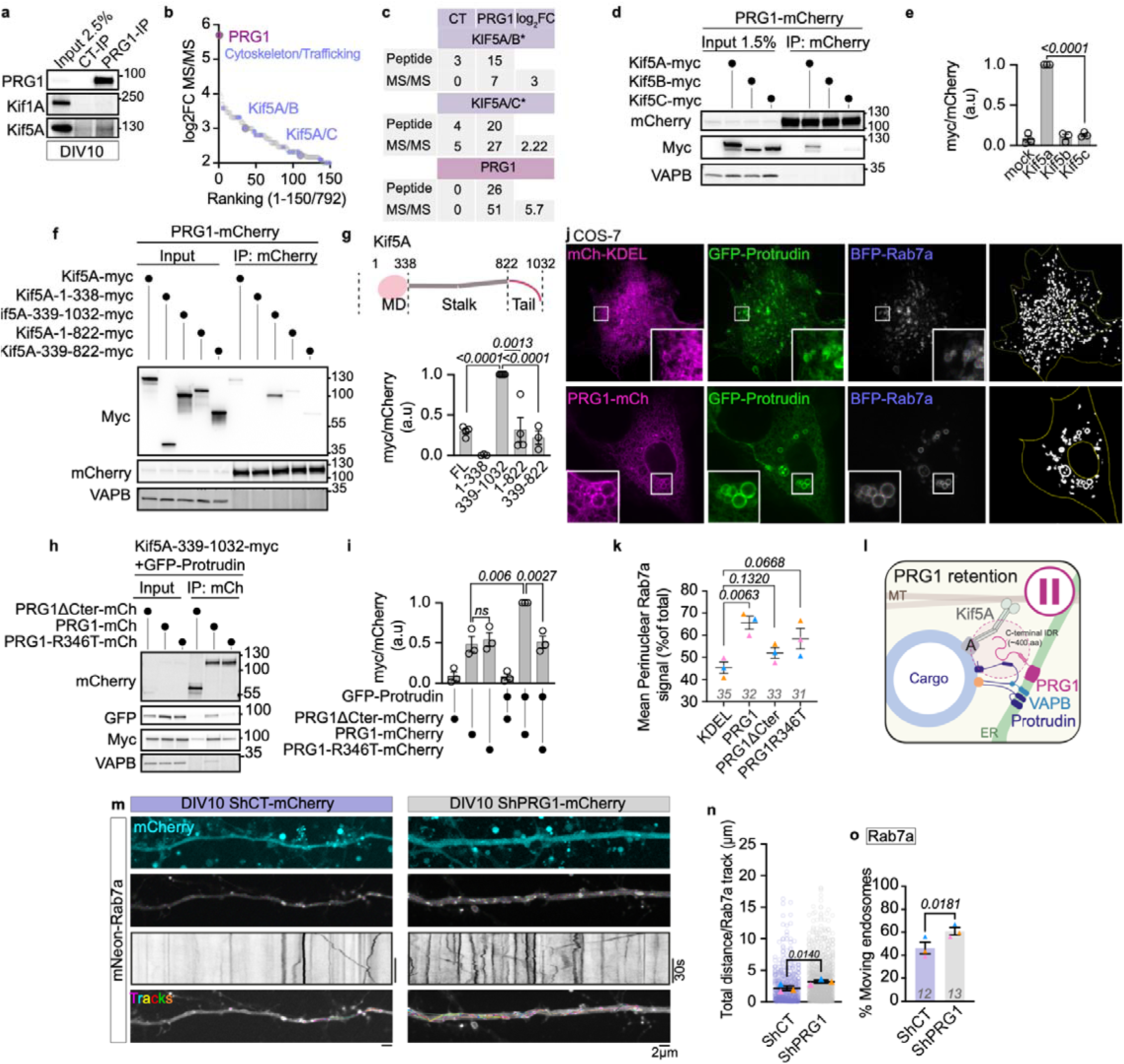
PRG1 complexes with KIF5A and Protrudin to pause late endosome trafficking. (a) Immunoblot analysis of endogenous PRG1 immunoprecipitation compared with a control IgG from DIV10 hippocampal neurons, showing co-immunoprecipitation of KIF5A, but not KIF1A. Input represents 2.5% of the total lysate used for immunoprecipitation. (b) Ranked proteins identified by mass spectrometry following endogenous PRG1 immunoprecipitation from DIV10 hippocampal neurons. PRG1, KIF5A/B/C and proteins associated with cytoskeletal organization and trafficking are highlighted. (c) Summary of peptide counts and MS/MS spectral counts for KIF5A/B/C and PRG1 identified by mass spectrometry following PRG1 immunoprecipitation compared with a control IgG. (d) Immunoblot analysis of mCherry immunoprecipitation from HEK293T cells expressing PRG1-mCherry together with KIF5A-myc, KIF5B-myc or KIF5C-myc. Input represents 1.5% of the total lysate used for immunoprecipitation. (e) Quantification of co-immunoprecipitated myc signal normalized to immunoprecipitated mCherry signal, showing a preferential interaction between PRG1 and KIF5A compared with KIF5B or KIF5C. Values were normalized to the maximum within each experiment and are presented as mean ± SEM from n=3 independent experiments compared using unpaired t-tests. (f) Immunoblot analysis of mCherry immunoprecipitation from HEK293T cells co-expressing PRG1-mCherry and the indicated KIF5A-myc constructs. Input represents 1.5% of the total lysate used for immunoprecipitation. (g) Schematic representation of KIF5A constructs used to identify the region involved in PRG1 interaction. Quantification of co-immunoprecipitated myc signal normalized to mCherry, showing that PRG1 preferentially interacts with the C-terminal region of KIF5A. Values were normalized to the maximum within each experiment and are presented as mean ± SEM from at least 3 independent experiments compared using unpaired t-tests. (h) Immunoblot analysis of mCherry immunoprecipitation from HEK293T cells expressing PRG1-mCherry, PRG1-R346T-mCherry or PRG1-ΔCter-mCherry together with KIF5A-339–1032-myc, in the presence of GFP-Protrudin. Input represents 1.5% of the total lysate used for immunoprecipitation. (i) Quantification of (h) and Fig. S4a, co-immunoprecipitated myc signal normalized to mCherry, showing enhanced PRG1-mCherry/KIF5A-339-1032-myc association in the presence of GFP-Protrudin and stronger co-immunoprecipitation with wild-type PRG1 compared with the indicated PRG1 mutants. Values were normalized to the maximum within each experiment and are presented as mean ± SEM from n=3 independent experiments compared using unpaired t-tests. (j) Representative images of COS-7 cells expressing mCherry-KDEL (ER), GFP-Protrudin and BFP-Rab7a (top panels), or PRG1-mCherry, GFP-Protrudin and BFP-Rab7a (bottom panels), illustrating changes in the intracellular distribution of Rab7a-positive endosomes. Magnified views of the boxed regions show the localization of Protrudin and PRG1 on Rab7a-positive endosomes. Cell outlines and segmented endosome masks used for quantification are shown on the right. (k) Quantification of perinuclear Rab7a localization in cells expressing mCherry-KDEL, PRG1-mCherry, PRG1ΔCter-mCherry or PRG1-R346T-mCherry. The perinuclear region was defined as the cytoplasmic area within 8 µm of the nuclear outline. Rab7a-positive pixels within this region were expressed as a percentage of all Rab7a-positive pixels in the cytoplasm. Each triangle represents the mean of an independent experiment (n = 3), 31 to 35 cells were analyzed per condition. Bars indicate mean ± SEM. P values were calculated using unpaired t-tests. (l) Proposed model of PRG1-mediated vesicle retention at ER-late endosome-microtubule interfaces through its association with KIF5A and the ER-endosome contact site proteins Protrudin and VAPB. Both the C-terminal intrinsically disordered region (IDR) and the amphipathic helix of PRG1 are necessary to form the complex. (m) Representative images, kymographs and TrackMate-generated tracks from DIV10 hippocampal neurons expressing mNeon-Rab7a under control or shPRG1 knockdown conditions. The shRNA-mCherry cassette is shown in cyan. (n) Quantification of the total distance traveled by Rab7a-positive endosomes in control and PRG1-depleted neurons. Data are mean ± SEM from at least 3 independent experiments compared using paired t-tests; 12-14 neurons analyzed. (o) Quantification of the mobile fraction of Rab7a-positive endosomes in control and PRG1-depleted neurons. Data are mean ± SEM from at least 3 independent experiments compared using paired t-tests; the number of neurons analyzed is indicated on each graph.

We wondered if PRG1 competes with Protrudin for KIF5A binding. To test this, we co-expressed PRG1-mCherry with “open” KIF5A-339-1032-myc with and without GFP-Protrudin in HEK293T cells. PRG1-mCherry was purified on mCherry-trap beads as before. Surprisingly, co-expression of Protrudin led to an increase in KIF5A-339-1032-myc binding. Notably, Protrudin and VAPB were also co-immunoprecipitated with PRG1 (Fig. 5h,i and S4a). To investigate the specificity and functional importance of this interaction, we compared whether KIF5A, Protrudin, and VAPB would also be co-purified by the PRG1 mutants that do not attenuate trafficking: PRG1-ΔCter and PRG1-R346T. Importantly, these functionally impaired PRG1-mutants had a reduced capacity to co-precipitate Protrudin, and VAPB with KIF5A-339-1032 (Fig. 5h,i). We therefore propose a model in which PRG1, Protrudin, and VAPB can be found in complex together with KIF5A (Fig. 5l).

To further investigate whether PRG1 could modulate Protrudin regulated LE trafficking, we co-transfected COS-7 cells with GFP-Protrudin, a LE marker (BFP-Rab7a) and with either a general ER marker (mCherry-KDEL) or with PRG1-mCherry or PRG1 mutants. LEs were wrapped by the ER and distributed throughout the cytoplasm in cells expressing Protrudin and mCherry-KDEL. In cells expressing Protrudin and PRG1, LEs were still wrapped with ER but strikingly clustered in the perinuclear region (Fig. 5j,k). Cells expressing PRG1-ΔCter and PRG1-R346T displayed intermediate phenotypes (Fig. 5k and Fig. S4b). As expected, Protrudin and PRG1 co-expression did not wrap or redistribute Rab5-positive early endosomes (Fig. S4c).

Protrudin and PRG1 are both expressed in developing neurons although only PRG1 levels increase between DIV5 and DIV10 (Fig. S1h,i and Fig. S4h,i). We therefore asked whether LE trafficking is also altered by PRG1 levels in DIV10 neurons. We performed PRG1 shRNA depletion experiments as before and scored the dynamics of Rab7a-positive LEs. PRG1 depletion significantly increased the distance traveled and the mobile fraction of Rab7a-endosomes (Fig. 5m-o). For comparison, PRG1 depletion did not increase trafficking of Rab5-positive early endosomes (Fig. S4d-f). Notably, PRG1 depletion did not alter Protrudin expression at DIV10 (Fig. S4g). These data support a role for PRG1 in restricting selective cargo mobility across different cellular contexts. Together, our data suggest that PRG1 could attenuate trafficking at a specific developmental stage by a stalled KIF5A contact site complex. At ER-LE contact sites, this complex is likely to include the Protrudin/VAPB machinery. Whether PRG1 uses distinct ER contact site machineries to pause KIF5A at SVP and DCV contacts remains to be determined.

### Cryo-ET reveals PRG1-dependent remodeling of ER-vesicle-microtubule interfaces

So far, our data suggest that PRG1 might promote the assembly of a stalled motor complex at ER contact sites. We performed cryo-electron tomography to gain insight into the three-dimensional (3D) spatial organization of ER contact sites in hippocampal neurons at DIV5 and DIV10, and at DIV10 in PRG1-depleted neurons. Western Blot, qPCR and fluorescence imaging combined with DIC confirmed efficient PRG1 depletion (Fig. S5a-d). Neurons were cultured on gold EM grids and plunge-frozen for cryo-electron tomography, allowing organelles and microtubules (MTs) to be resolved at nanometer resolution, as in (Zamponi et al., 2022)(Fig. 6a-c). 3D reconstructions also resolved axonal ER ladder structures (magenta; Fig. 6a-c), whose organization appeared more prominent at DIV5 than at DIV10, as predicted (Zamponi et al., 2022). Axonal vesicles were classified by diameter as SVs or SVPs (40-70 nm, yellow) and DCVs (70-155 nm, orange) (Foster et al., 2022; Takamori et al., 2006). Vesicle diameter distributions were comparable between conditions (Fig. S5e,f). Across three independent experiments and at least 8 tomograms per condition, we modeled vesicles that were closely apposed to the ER and could visualize clear examples of electron-dense, tether-like structures between the ER and these vesicles for a significant percentage of vesicles modeled (14.4% of 111 vesicles for DIV5, 13.1% of 199 vesicles for DIV10, 9.1% of 274 vesicles for DIV10 shCT and 13.1% of 252 vesicles in DIV10 shPRG1 tomograms). These data provide the strongest evidence that complexes are present between these organelles (Fig. 6d, S5i and movie S5). Observed tethers ranged from 6.4 to 55.8 nm in length and displayed irregular and folded morphologies; tether length distribution remained comparable across developmental stages and following PRG1 depletion (Fig. 6d,f and S5g,h). Therefore, tethers remain in PRG1-depleted neurons (Fig. 6d,f).

**Figure 6.**
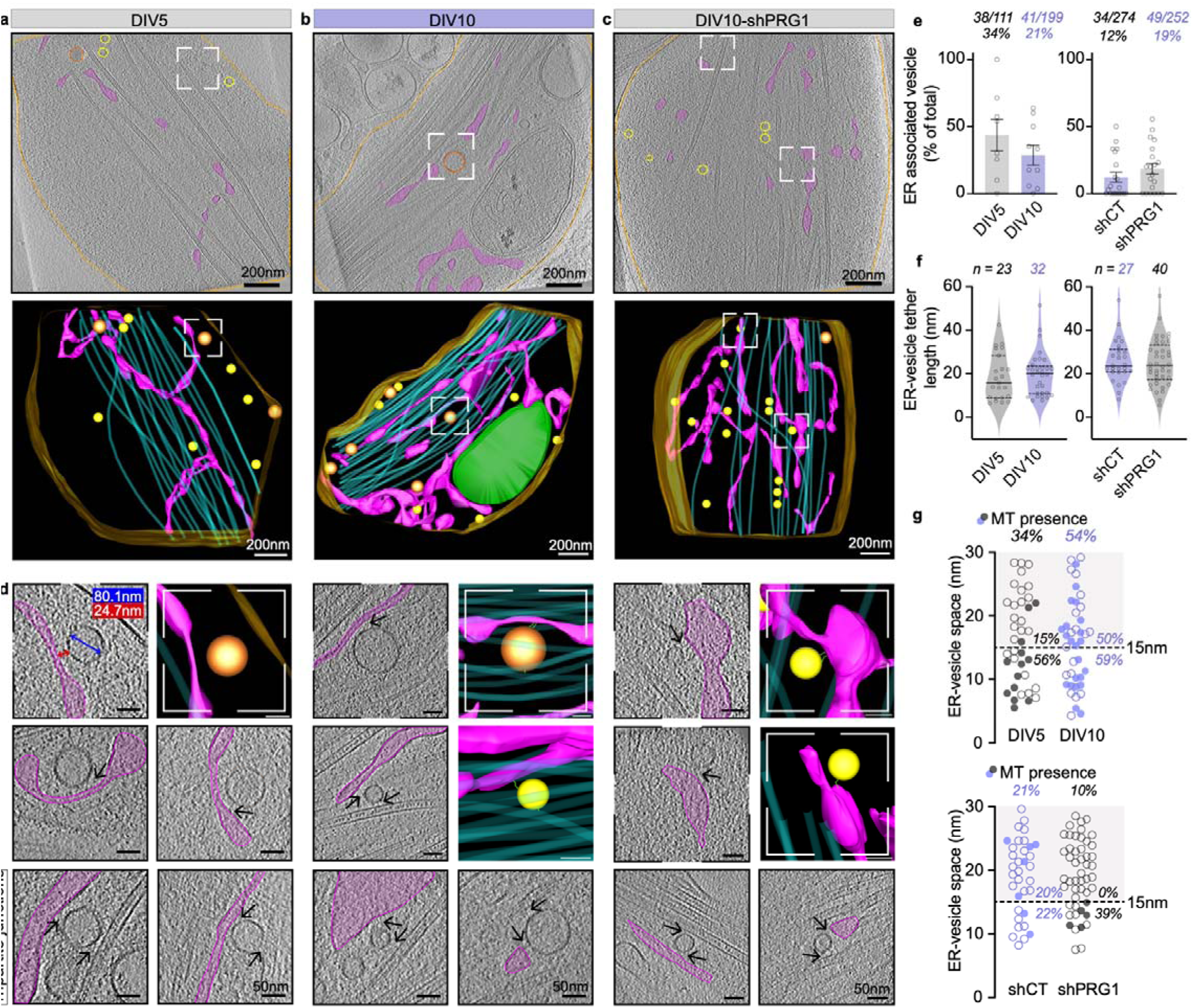
Cryo-ET reveals PRG1-dependent remodeling of ER-vesicle-microtubule interfaces. (a,b,c) Tomograms and corresponding 3D models of axonal ER (pink), MTs (cyan), plasma membrane (gold), mitochondria (green), SVs/SVPs (yellow, based on diameter range 40-70 nm) and DCVs (orange, 70-155 nm) imaged by cryo-EM in DIV5 (a), DIV10 (b) and DIV10 PRG1 knockdown hippocampal neurons (c). The cropped and enlarged area highlights a vesicle that is in close proximity to the ER. (d) Enlarged views illustrating ER-vesicle contacts and tripartite ER-vesicle-MT junctions. Blue and red arrows indicate vesicle diameter and ER-vesicle membrane distance, respectively. Black arrows indicate densities consistent with physical tethers between vesicles and the ER and/or MTs. (e) Percentage of vesicles associated with the ER in DIV5 versus DIV10 neurons and DIV10 shControl versus shPRG1 neurons. ER association was defined as an ER-vesicle membrane distance <30 nm, regardless of tethering. Each point represents one tomogram; bars show the mean ± SEM across tomograms (DIV5 n=8; DIV10 n=10; shCT n=20; shPRG1 n=20 tomograms from three independent experiments). ER-associated vesicles over the total number of vesicles and the corresponding percentages are indicated at the top of the graphs. (f) Distributions of individual ER-vesicle tether lengths in the indicated conditions, pooled across experiments. Each point represents one measured tether; multiple tethers associated with the same vesicle were plotted separately. Sample sizes are indicated above each distribution. (g) ER-vesicle distance for individual vesicles located <30 nm from the ER, comparing DIV5 with DIV10 neurons (top) and DIV10 shControl with shPRG1 neurons (bottom). Each point represents one vesicle; filled circles indicate that the vesicle was also located <30 nm from at least one MT, highlighting tripartite ER-vesicle-MT junctions. Percentages above each condition indicate the proportion of ER-associated vesicles also associated with MTs. The dashed line marks a separation of contacts in two size categories; percentages within each distance interval indicate the proportion of vesicles also associated with MTs among ER contacts at 0 to 15 nm or 15 to 30 nm.

Because the ability to visualize individual tethers in cryo-ET depends on their orientation and can be compromised by missing wedge artifacts (Y. T. Liu et al., 2022), we believe these percents may underestimate the actual number of tethered vesicles. We therefore complemented this analysis with a proximity-based metric, quantifying ER-associated vesicles as those with an ER-vesicle membrane separation of <30nm. This strategy also has limitations but has been an acceptable standard for scoring contacts in published works (Friedman et al., 2013; Scorrano et al., 2019)(Fig. 6e). For each of these ER-associated vesicles (38 vesicles at DIV5, 41 at DIV10, 34 in shCT and 49 in shPRG1 neurons), we scored whether the vesicle was also associated with a MT (within 30nm) indicative of a tripartite junction between the ER, vesicle, and MT. ER-associated vesicles at DIV10 were more likely than DIV5 vesicles to be positioned near one or multiple MTs, increasing from 34% to 54% (Fig. 6g, top), suggesting a developmental remodeling of ER-vesicle-MT junctions. The proportion of ER-MT junctions also decreased following PRG1-depletion from 21% to 10% (Fig. 6g, bottom). Interestingly, no tripartite junctions were detected within the 15-30 nm ER-vesicle distance range after PRG1-depletion, mirroring the pronounced developmental increase in this specific fraction from DIV5 to DIV10 (from 15 to 50%) (Fig. 6g, top). Although these Cryo-ET proximity-based measurements provide only limited evidence of structural remodeling, these observations suggest that PRG1 is not essential for establishing ER-vesicle contacts but may contribute to organizing a vesicle-ER-MT tripartite junction and support the model whereby PRG1 promotes vesicle retention at these sites through its association with KIF5A and ER contact-site machinery.

### PRG1 loss increases synapse assembly and hyperexcitability at DIV10

Our experiments highlight PRG1 as a necessary contributor to the maturation-associated modulation of axonal vesicle trafficking. We therefore wondered whether PRG1-dependent modulation of vesicle trafficking could influence the timing of synapse formation during early development. To address this question, rat hippocampal neurons were infected with a PRG1-targeting shRNA (shPRG1-mCherry) or control (shCT-mCherry) at DIV4 to deplete PRG1 and synapse formation was analyzed at DIV10. Putative synapses were visualized by performing immunofluorescence labeling against the presynaptic protein Bassoon and the postsynaptic protein Homer (as described previously (Verstraelen et al., 2020)). Synapses were defined as elongated puncta (≥ 0.2 µm) where Bassoon and Homer signals partially overlapped along neuronal processes; the quantification method for synapse density and size is illustrated in Fig. 7a. PRG1 depletion (by shRNA) resulted in a 2.6-fold increase in the number of synapse densities compared with the shCT controls (Fig. 7b-d). In addition, synaptic puncta measured in PRG1-depleted neurons were significantly larger, with a greater proportion exceeding 0.5 µm (Fig. 7e); consistent with increased synaptic maturation at DIV10 (Wittenmayer et al., 2009).

**Figure 7.**
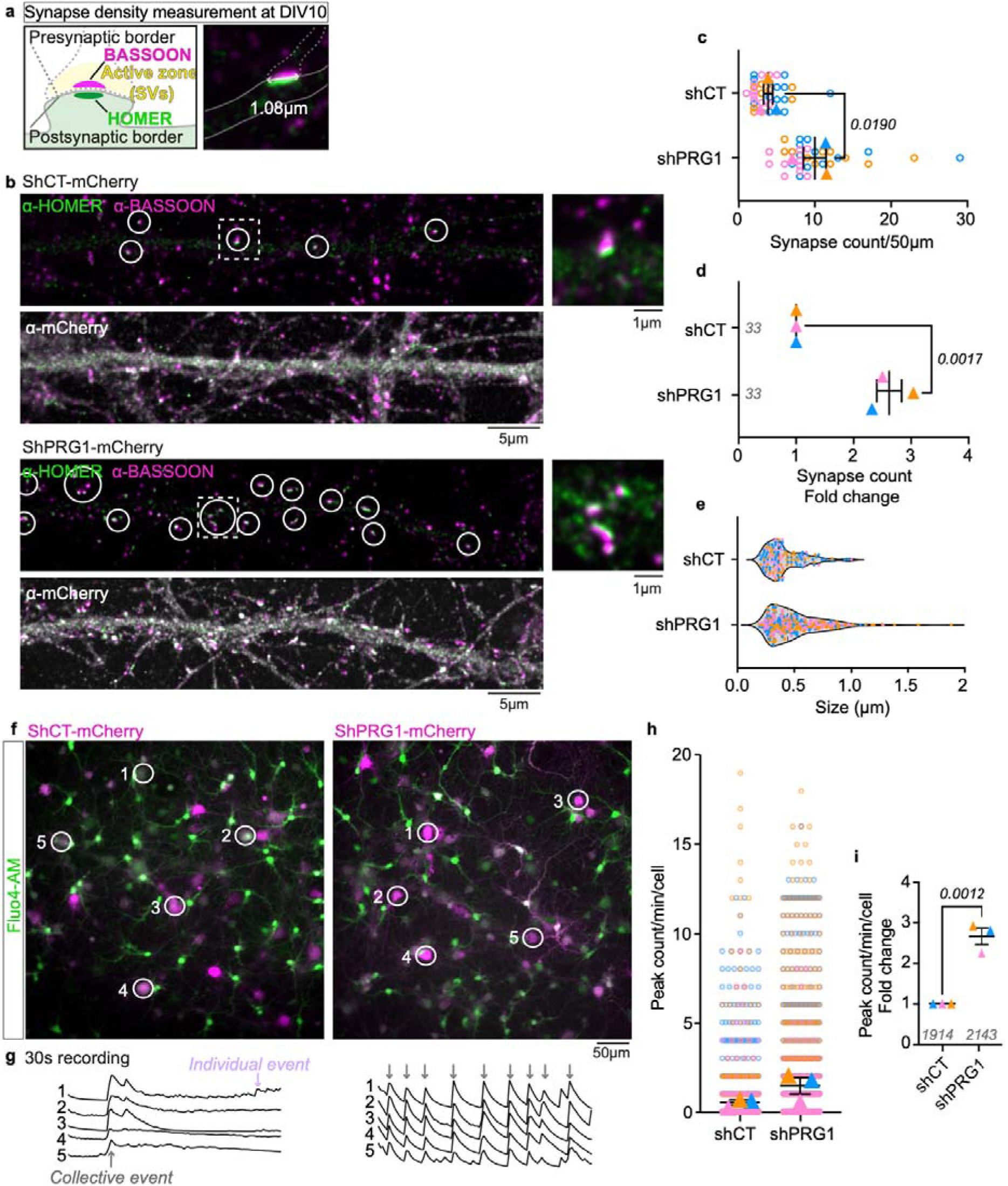
PRG1 loss increases synapse assembly and hyperexcitability at DIV10. (a) Schematic illustrating the quantification strategy used to measure putative synapse density and size based on Bassoon/Homer pattern and pixel colocalization. (b) Representative immunofluorescence images and cropped and zoomed areas of the presynaptic marker Bassoon (magenta) and the postsynaptic marker Homer (green) in DIV10 rat hippocampal neurons expressing control shRNA-mCherry (shCT) or PRG1-targeting shRNA-mCherry (shPRG1). An antibody against mCherry (grey) was used to identify the shRNA-infected cells. (c,d) Quantification of putative synapse density data from experiment (b) revealed a ∼2.6-fold increase in Bassoon/Homer-positive synapses in PRG1-KD neurons compared with controls. Data are mean ± SEM from n=3 independent experiments compared through unpaired t-tests, 33 neurons were analyzed in total per condition. (e) Distribution of synapse size in control and PRG1-KD neurons, revealing a higher proportion of large synapses (>0.5 µm) in PRG1-deficient conditions. (f,g) Calcium imaging using the Fluo4-AM probe (green) in live developing neuronal cultures imaged at DIV10. Representative images (f) and fluorescence traces (g) illustrate increased frequency and synchrony of spontaneous calcium events in PRG1-KD (magenta) neurons compared with controls. Recordings were acquired over 1 min at 3 frames s⁻ ¹. (h,i) Quantification of network-synchronized calcium event frequency in shRNA-mCherry positive cells, showing a ∼2.7-fold increase in PRG1-KD neurons. Data are mean ± SEM from n=3 independent experiments compared with an unpaired t-test, 12 fields per condition, ∼ 150 neurons per field.

Finally, we measured whether the enhanced synapse formation observed upon PRG1 depletion translates into a higher neuronal activity (Matz et al., 2010). PRG1 was depleted by shRNA (shPRG1-mCherry) or control (shCT-mCherry) as previously described. Neuronal activity was assessed at DIV10 by calcium imaging: rat hippocampal neurons were loaded with the fluorescent Ca^2+^ probe Fluo-4 AM, and spontaneous Ca^2+^ events were recorded over 1 minute at 3 frames per second in an artificial cerebrospinal fluid solution (Bardy et al., 2015), as a readout of neuronal network activity (Grienberger & Konnerth, 2012; Radotić et al., 2018). Control neurons at DIV10 showed mostly isolated single-cell events, whereas PRG1-KD neurons displayed frequent collective bursts and faster rhythmic oscillations (Fig. 7f,g and movie S6). The frequency of collective events was quantified, and PRG1-deficient neurons exhibited nearly three times more network-synchronized events than control neurons (Fig. 7h,i). These observations indicate that the loss of PRG1 promotes coordinated neuronal activity, consistent with enhanced synapse maturation and epileptic seizures or hyperexcitability observed *in vivo* (Knierim et al., 2023; Trimbuch et al., 2009; Vogt et al., 2016).

## Discussion

Here, we provide the first characterization of membrane contact sites between the axonal ER and SVPs or DCVs. We identify PRG1 as an ER-localized factor capable of pausing axonal vesicle mobility. PRG1 expression increases during a defined developmental window and coincides with a transient slowdown of vesicle transport around DIV10, a critical period of active synapse formation (Grabrucker et al., 2009; Verstraelen et al., 2020). Gain-and loss-of-function experiments indicate that PRG1 levels influence vesicle mobility: PRG1 overexpression restrains trafficking, whereas PRG1 depletion restores high vesicle mobility and is accompanied by enhanced synaptic assembly and neuronal activity at DIV10. The similar expression profile of PRG1 in humans, together with the loss of trafficking regulation caused by a patient-associated epilepsy mutation, indicate that this function may be relevant to human neuronal development. Together, these findings suggest that dysregulated vesicle transport may facilitate the delivery and accumulation of presynaptic material at nascent synapses (Ahmari et al., 2000; Rizalar et al., 2023; Shapira et al., 2003; Vukoja et al., 2018), thereby promoting synaptic maturation and contributing to network hyperexcitability, characteristic of juvenile epileptic disorders associated with PRG1 dysfunction (Knierim et al., 2023).

Mechanistically, we found that PRG1 promotes vesicle pausing at tripartite ER-vesicle-MT junctions. Indeed, PRG1 associates with the motor KIF5A and the Protrudin-VAPB machinery at ER-LE contact sites, providing a molecular basis for its involvement in LE trafficking regulation. Although PRG1 regulates the mobility of Syt1-Syt4- and Rab7a-positive cargo populations, the molecular partners allowing SVP and DCV trafficking control remain unidentified. Whether PRG1 relies on a similar contact-site machinery or distinct cargo-specific adaptors will be an important question for future studies. Importantly, while KIF1A is considered a major motor for SVP and DCV transport (Lo et al., 2011; Okada et al., 1995; Stucchi et al., 2018), the heterogeneity of presynaptic precursor populations during development (Petzoldt, 2023) together with the identification of kinesin-1 motors in purified SV fractions (Takamori et al., 2006), raises the possibility that different kinesin/dynein motors can coexist on the same cargo. Thus, PRG1 could selectively retain a cargo primarily transported by KIF1A, KIF5A, or dynein, by interacting with additional KIF5A molecules present. Indeed, perturbation of one motor system has been shown to impair both transport directions (Anazawa et al., 2022; J. S. Liu et al., 2012). Consistent with this, PRG1 depletion increases both anterograde and retrograde mobility of SVPs, DCVs, and Rab7a-late endosomes without altering their maximum velocity.

Remarkably, the disease-associated R346T mutation within PRG1’s amphipathic helix (AH) disrupts its association with Protrudin-VAPB and abolishes its trafficking regulatory function, despite preserving KIF5A association. These data indicate that motor association alone is not sufficient for PRG1 function. Whether the AH directly mediates interaction with partners such as Protrudin for PRG1 to be properly recruited in contact sites or provides membrane anchoring required for the complex assembly remains unexplored. Interestingly, PRG2, another PRG family member, also contains a cytosolic AH with markedly reduced amphipathicity (hydrophobic moment μH = 0.239 versus 0.556 for PRG1 AH)(Fuchs et al., 2022; Gautier et al., 2008) and also modulates SVP mobility upon overexpression, albeit to a lesser extent than PRG1 (Fig. S6a-c). This suggests that PRG family members may constitute a molecular toolkit to tune vesicular trafficking throughout neuronal development with distinct mechanisms and efficiencies.

Intracellular calcium signaling is a key regulator of vesicle trafficking and synaptic function (Südhof, 2012; Xue et al., 2021). An important question is how PRG1 activity is regulated beyond changes in its expression during neuronal development. Notably, PRG1 contains a C-terminal Calmodulin/Ca^2+^ binding region (Tokumitsu et al., 2010), raising the possibility that its trafficking regulatory function is sensitive to local calcium signals. Indeed, Ca^2+^/Calmodulin has also been implicated in regulating the SNARE complex conformation (de Haro et al., 2004), Kif1a-driven DCV transport (Stucchi et al., 2018) and more recently, in the interaction with VPS13C, a lipid transfer protein at ER-lysosome MCSs (Li et al., 2026). Moreover, PRG1 associates with PPP2R2A, a regulatory subunit of the protein phosphatase 2a (PP2A)(X. Liu et al., 2016; Vogt et al., 2016). Whether PRG1-dependent trafficking regulation, or its association with KIF5A is gated by phosphorylation or by Calmodulin/Ca^2+^ signaling remains to be determined.

A key conceptual implication of our work is that the transient reduction in axonal vesicle transport represents a regulatory stage during neuronal circuit assembly. We propose that developmental PRG1 expression at the ER contributes to temporal coordination between vesicle delivery, synaptic assembly, and the emergence of network activity. Beyond this temporal role, our findings raise a broader question: does the developmentally regulated architecture of the ER and its associated MCSs in the growing axon also contribute to the spatial organization of presynaptic sites? More specifically, does the repetitive spacing of the axonal ER ladder (1-2 µm) provide the initial roadmap for the (∼2–5 µm) spacing of presynaptic compartments (Mishchenko et al., 2010; Shepherd & Harris, 1998; Takács et al., 2018)?

---

**Figure S1.**
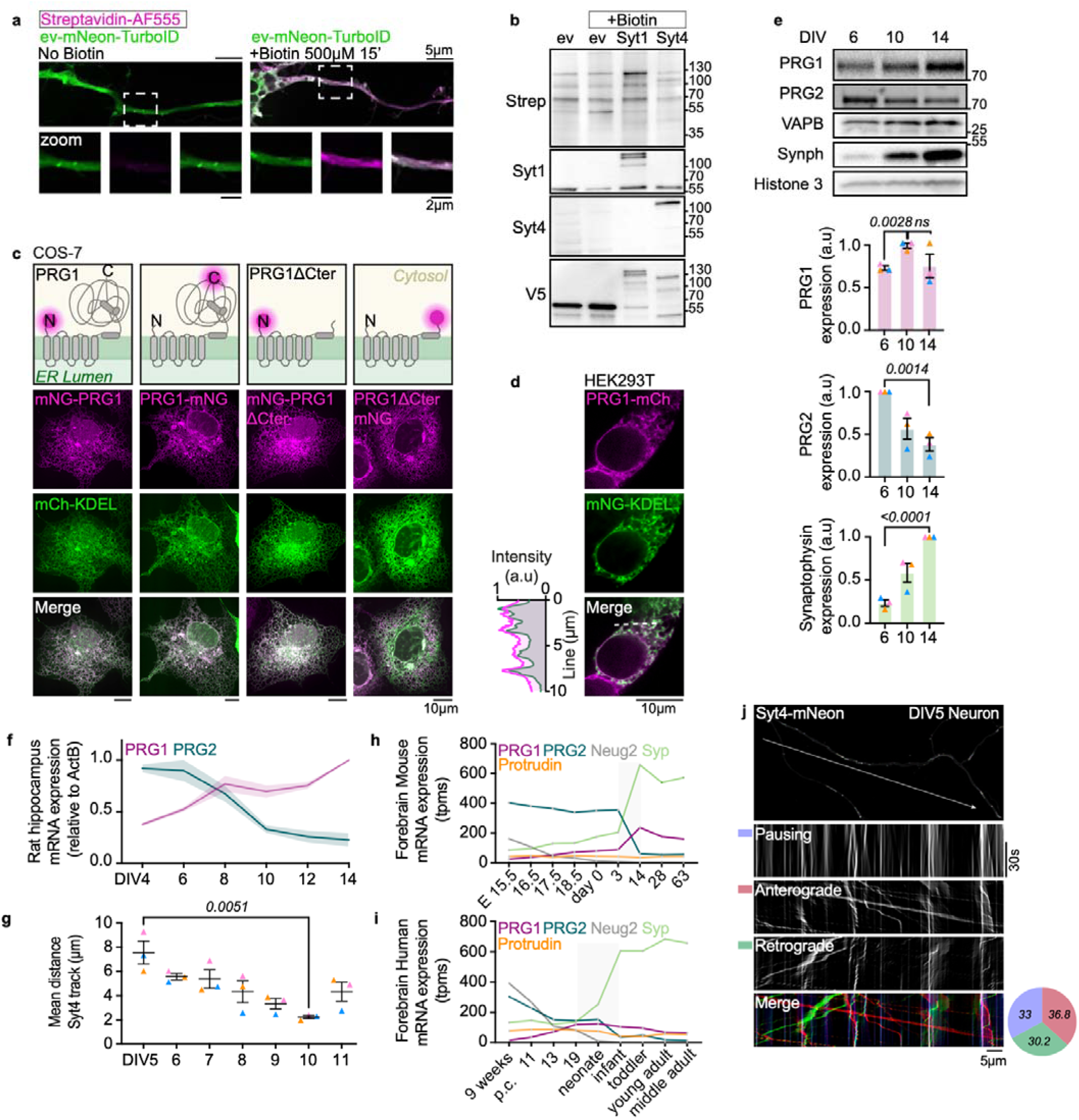
related to Figure 2 (a) Representative images showing biotinylated proteins detected by streptavidin-AF555 staining in hippocampal neurons expressing the control empty-vector (EV-mNeon-V5-TurboID) before and after biotin incubation. The cytosolic streptavidin signal increased after treatment. (b) Immunoblot analysis of TurboID-mediated biotinylation in neurons expressing Syt1-mNeon-V5-TurboID, Syt4-mNeon-V5-TurboID, or EV-mNeon-V5-TurboID, confirming specific protein biotinylation after treatment. (c) Predicted topology of PRG1 within the ER membrane based on AlphaFold, DISOPRED and TMHMM analyses and representative images of PRG1-mNeonGreen expressed in COS-7 cells, showing predominant localization to the endoplasmic reticulum, labeled with the KDEL marker. ER localization is preserved independently of tag orientation and upon deletion of PRG1 cytosolic domain. (d) Representative images of PRG1-mCherry expressed in HEK293T cells, together with corresponding line-scan intensity profiles across ER tubules, illustrating spatial overlap between PRG1-mCherry and the ER markers mNeonGreen-KDEL. (e) Immunoblot analyses of protein levels for Synaptophysin (Synph), PRG1, and PRG2 protein across developmental stages normalized to VAPB. Corresponding quantitative analyses of immunoblot data from 3 independent cultures. Data are mean ± SEM, unpaired t-tests. (f) Quantification of PRG1 and PRG2 mRNA levels in rat hippocampal neurons cultured from DIV4 to DIV14, showing a peak of PRG1 mRNA around DIV8 when PRG2 expression decreases. Data are mean ± SEM from n=4 independent cultures. (g) Experiment of Figure 2j,m shown here across developmental stages DIV5 to DIV11. Quantification of Syt4-mNeon labeled-vesicle total run distances indicate a progressive decrease in motility from DIV5 to DIV10, followed by a partial recovery. Data are mean ± SEM from n=3 independent experiments; 10 to 12 neurons per condition. Statistics: unpaired t-test. (h,i) Developmental expression profile of PRG1, PRG2, Synaptophysin (Syp), Neurogenin-2 (Neug2) and Protrudin mRNA in rodent (h) and human (i) forebrain extracted from publicly available brain expression atlases (E-MTAB-6814, Cardoso-Moreira etal., 2019) expressed in transcripts per million (tpms). (j) Representative Syt4-mNeon image and corresponding KymoClear analysis illustrating the classification of vesicle trafficking events as pausing, anterograde or retrograde. Kymographs were directionally filtered using KymoClear to separate each trafficking category, and the resulting events were quantified to determine their relative proportions. The merged image shows the three classified populations used to generate the pie charts in Figures 2 and 4. The arrow indicates the orientation of the axon.

**Figure S2.**
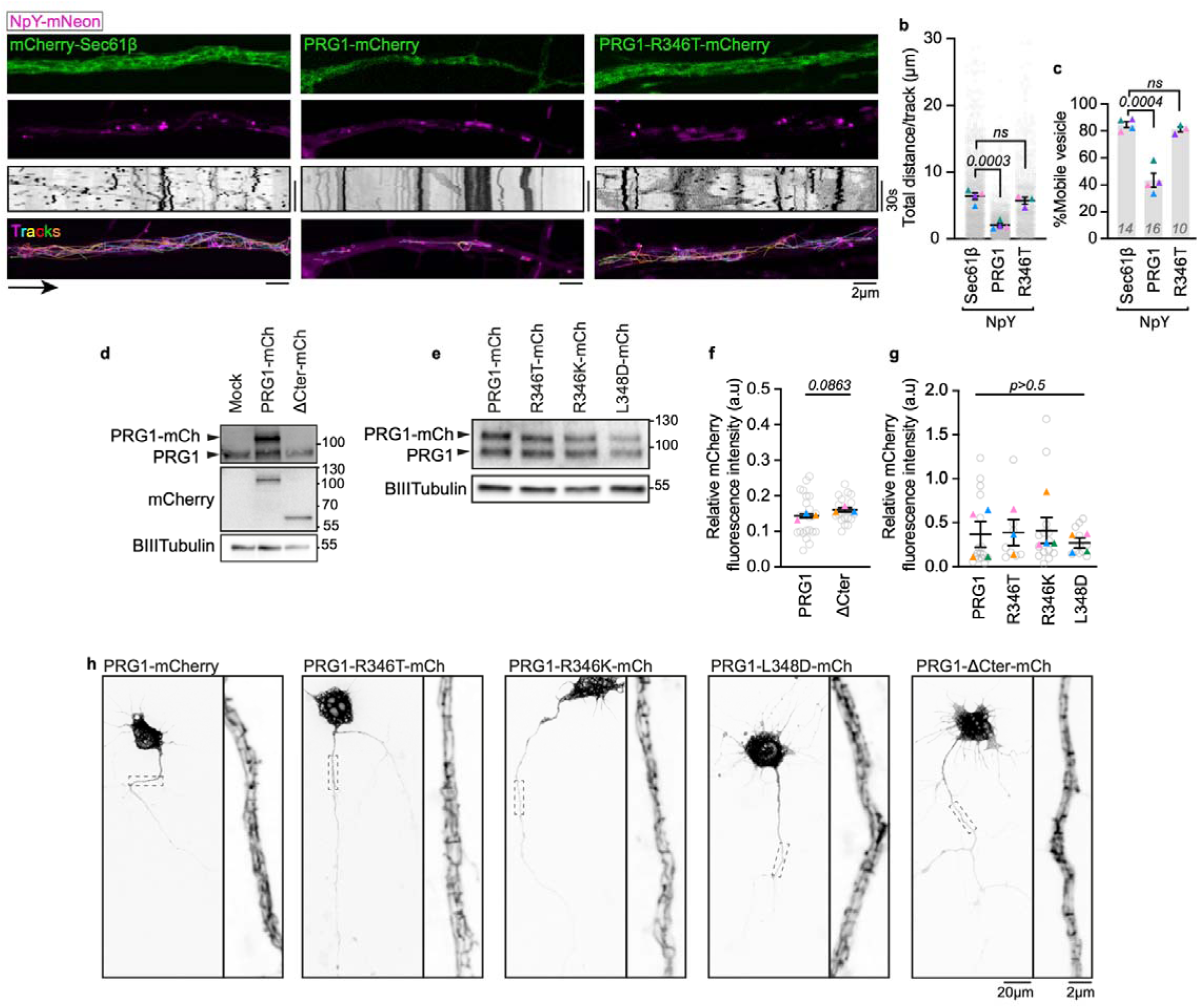
related to Figure 3 (a) Representative images, kymographs and TrackMate-generated tracks from 45 s movies of NpY-mNeon labeled vesicles (magenta) in DIV5 hippocampal neurons expressing mCherry-Sec61β as a control ER membrane protein, PRG1-mCherry or PRG1-R346T-mCherry (green). (b,c) Quantification of (b) run length and (c) mobile fraction of data from (a). Data are mean ± SEM from n=3 independent experiments compared through unpaired t-tests, 10 to 16 neurons were analyzed per condition. (d,e) Immunoblot analysis of PRG1-mCherry and the indicated PRG1 mutants overexpressed in DIV5 hippocampal neurons. βIII-Tubulin was used as a loading control. (f,g) Quantification of normalized PRG1-mCherry fluorescence intensity in the same neurons used for the vesicle-trafficking analyses in Fig. 3g and Fig. 3d, respectively, showing comparable expression levels between (f) PRG1 and PRG1-ΔCter and (g) PRG1 and the indicated amphipathic helix mutants. Data are mean ± SEM; n= 3 or 4 experiments. Statistics: (f) unpaired t-test and (g) one-way ANOVA. (h) Representative images of DIV5 hippocampal neurons expressing PRG1-mCherry or the indicated PRG1 mutants, showing overall neuronal morphology and distribution of the mCherry signal. Magnified views of the boxed axonal regions are shown on the right of each image.

**Figure S3.**
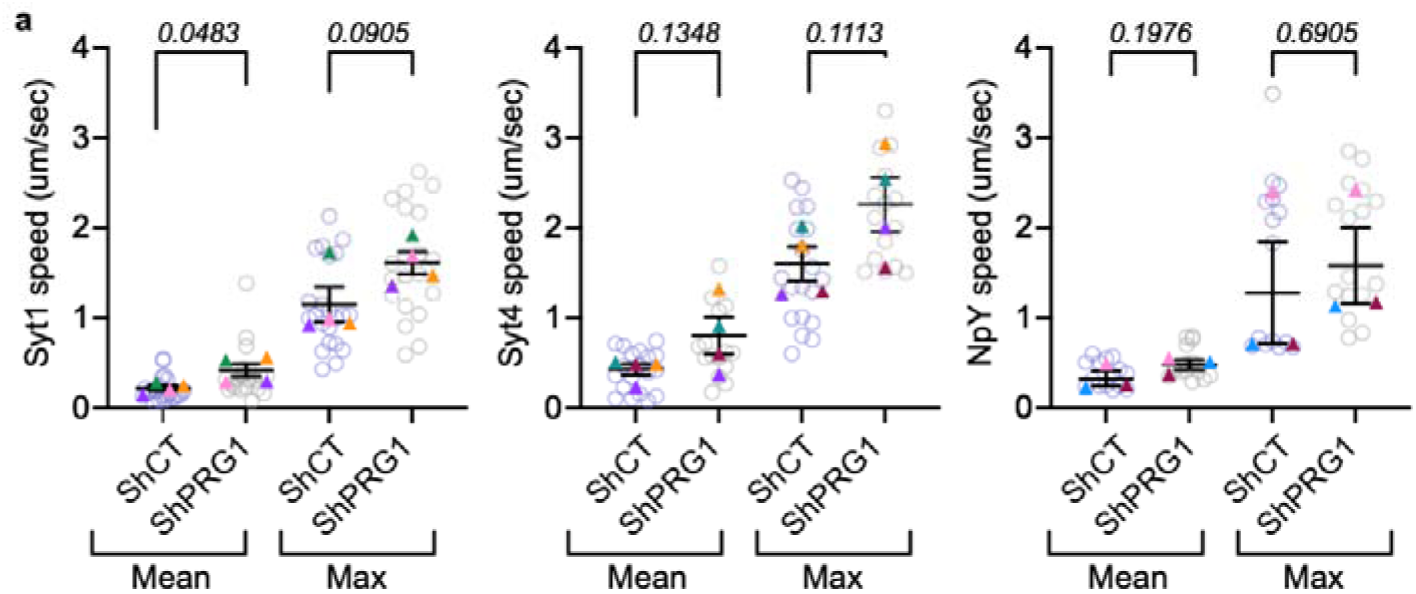
related to Figure 4 (a) Quantification of the mean and maximum speed of moving Syt1-mNeon-, Syt4-mNeon-and NpY-mNeon-labeled vesicles in control and PRG1-depleted DIV10 neurons. Only vesicles with a displacement greater than 1.5 µm were included in the analysis. Data are mean ± SEM from at least 3 independent experiments compared using unpaired t-tests.

**Figure S4.**
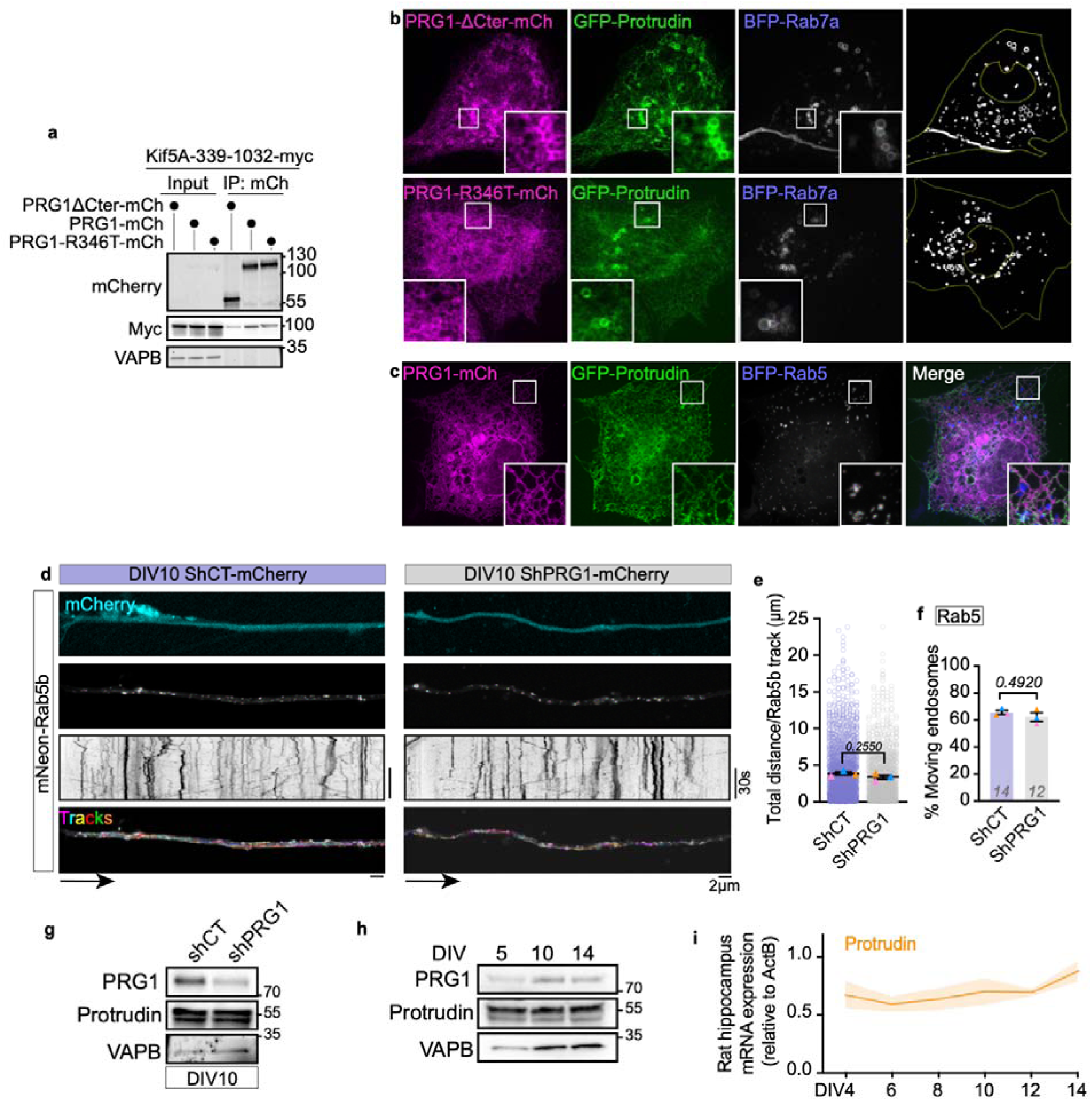
related to Figure 5 (a) Immunoblot analysis of mCherry immunoprecipitation from HEK293T cells expressing PRG1-mCherry, PRG1-R346T-mCherry or PRG1-ΔCter-mCherry together with KIF5A-339–1032-myc, in the absence of GFP-Protrudin. Input represents 1.5% of the total lysate used for immunoprecipitation. (b) Representative images of COS-7 cells expressing PRG1-ΔCter-mCherry or PRG1-R346T-mCherry together with GFP-Protrudin and BFP-Rab7a. Magnified views show the localization of Protrudin and PRG1 mutants on Rab7a-positive endosomes. Cell outlines and segmented endosome masks used for quantification are shown on the right. (c) Representative images of COS-7 cells expressing PRG1-mCherry, GFP-Protrudin and BFP-Rab5. Magnified views show the localization of PRG1 and Protrudin relative to Rab5-positive endosomes. (d) Representative images, kymographs and TrackMate-generated tracks from DIV10 hippocampal neurons expressing mNeon-Rab5 under control or shPRG1 knockdown conditions. The shRNA-mCherry cassette is shown in cyan. (e) Quantification of the total distance traveled by Rab5-positive endosomes in control and PRG1-depleted neurons. Data are mean ± SEM from n=3 independent experiments compared using paired t-tests; 12-14 neurons analyzed in total per condition. (f) Quantification of the mobile fraction of Rab5-positive endosomes in control and PRG1-depleted neurons. Data are mean ± SEM from n=3 independent experiments compared using paired t-tests; the number of neurons analyzed is indicated on each graph. (g) Immunoblot analysis of endogenous PRG1 and Protrudin protein levels in DIV10 hippocampal neurons under control or shPRG1 knockdown conditions. VAPB was used as a loading control. (h) Immunoblot analysis of endogenous PRG1 and Protrudin protein levels in hippocampal neurons at DIV5, DIV10 and DIV14. VAPB was used as a control. (i) Quantification of Protrudin mRNA levels in rat hippocampal neurons cultured from DIV4 to DIV14, showing that Protrudin expression is steady. Data are mean ± SEM from 4 independent cultures, the same samples as in Fig. S1f.

**Figure S5.**
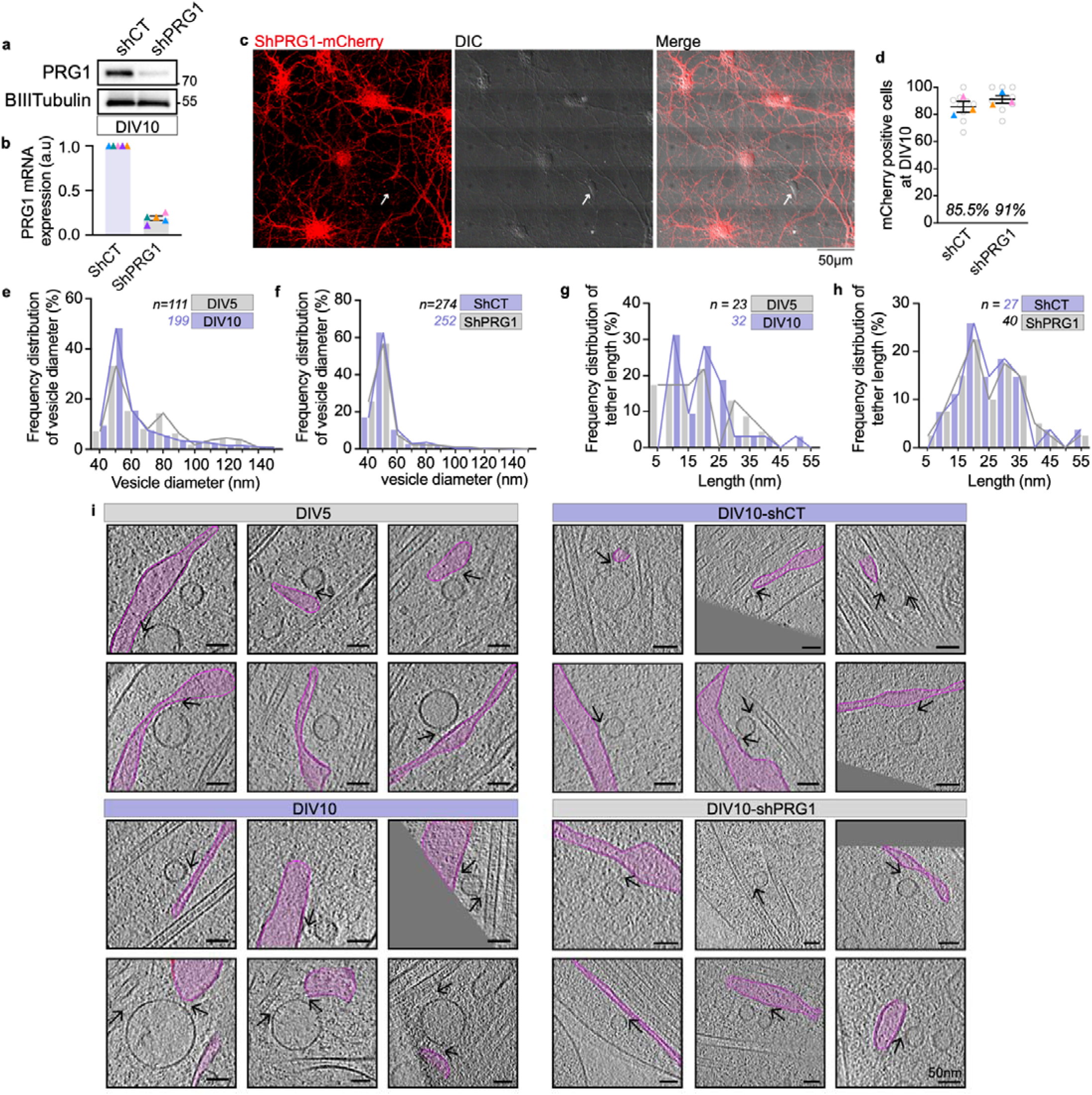
related to Figure 6 (a) Immunoblot analysis of endogenous PRG1 protein levels in DIV10 hippocampal neurons under control or shPRG1 knockdown conditions used for cryo-electron tomography. βIII-Tubulin was used as a loading control. (b) Quantification of PRG1 mRNA expression in control and PRG1-depleted neurons, normalized to shCT controls. Data are mean ± SEM from 5 independent experiments, 3 were used for cryo-electron tomography. (c) Representative fluorescence and DIC images of DIV10 hippocampal neurons expressing the shPRG1-mCherry cassette, illustrating neuronal morphology and the identification of transduced neurons used for cryo-electron tomography. The arrow indicates a non-transduced cell. (d) Quantification of the percentage of mCherry-positive neurons under shCT and shPRG1 conditions, showing comparable transduction efficiencies (85.5% and 91%, respectively). Data are mean ± SEM; n=3 individual experiments, 12 fields per condition. (e,f) Frequency distributions of vesicle diameters in DIV5 versus DIV10 hippocampal neurons (e) and DIV10 shControl versus shPRG1 neurons (f). All measured vesicles were included, irrespective of ER or MT association. Distributions were pooled across experiments and normalized to the total number of vesicles in each condition. Sample sizes are indicated. (g,h) Frequency distributions of individual ER-vesicle tether lengths in DIV5 versus DIV10 neurons (g) and DIV10 shControl versus shPRG1 neurons (h). Each measured tether was counted separately, including multiple tethers associated with the same vesicle. Distributions were pooled across experiments and normalized to the total number of measured tethers in each condition. Sample sizes are indicated. (i) Additional representative tomographic slices illustrating ER-vesicle contacts and ER-vesicle-MT associations in the indicated conditions. The ER is highlighted in pink. Black arrows indicate densities consistent with physical tethers between vesicles and the ER and/or MTs.

**Figure S6.**
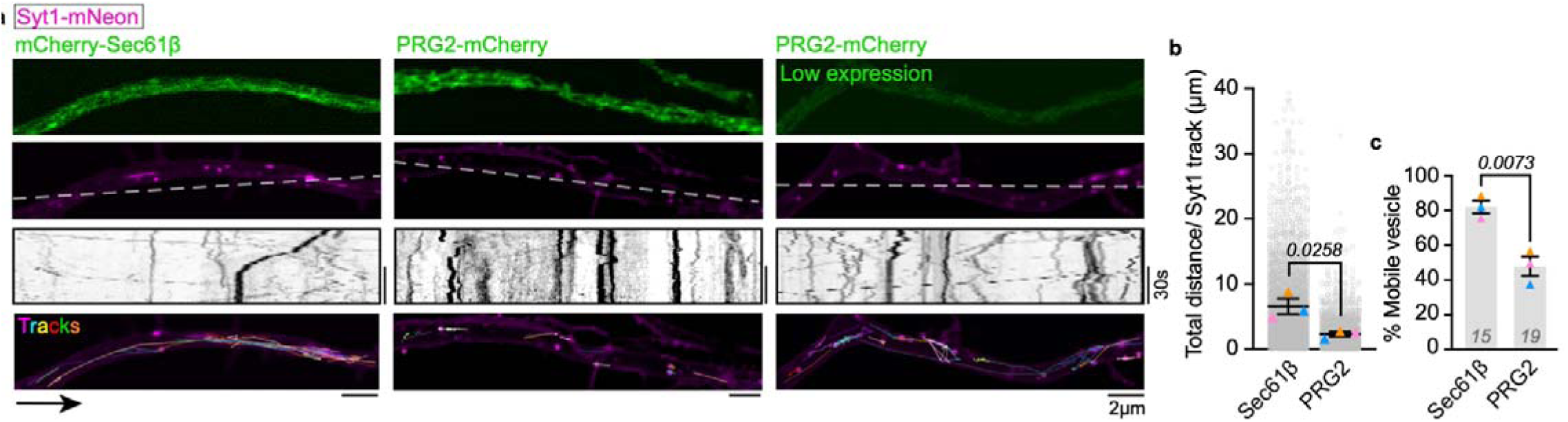
(a) Representative images, kymographs and TrackMate-generated tracks of Syt1-mNeon labeled vesicles (magenta) in DIV5 hippocampal neurons overexpressing mCherry-Sec61β or PRG2-mCherry at different expression levels (green), indicating that PRG2 also reduces axonal vesicle trafficking in rat hippocampal neurons in a concentration-dependent manner. (b,c) Quantification of (b) run length and (c) mobile fraction in neurons overexpressing PRG2-mCherry compared to mCherry-Sec61β. Data are mean ± SEM from n=3 independent experiments compared through unpaired t-tests, 15 to 19 neurons were analyzed in total per condition.

**Movie S1:** (Related to Fig. 1d-f) 1 min movie (∼0.5 frame s^-1^) of a DIV5 hippocampal rat neuron overexpressing the ER luminal mCherry-KDEL (green) and the DCV protein Syt4-mNeonGreen (magenta) showing vesicles trafficking alongside dynamic ER. White arrows on the cropped area indicate ER rungs where a vesicle stops or switches direction.

**Movie S2:** (Related to Fig. 2i) 45 s movie (∼0.9 frame s^-1^) of DIV5 or DIV10 hippocampal rat neurons overexpressing the SVP protein Syt1-mNeon (magenta) and the ER mCherry-KDEL marker (green) and accompanying TrackMate-generated tracks showing more intense trafficking at DIV5 and a rearrangement of ER shape and dynamics at later stages.

**Movie S3:** (Related to Fig. 3c) 45 s movie (∼0.7 frame s^-1^) of DIV5 hippocampal rat neurons overexpressing Syt1-mNeon (magenta) alongside the ER protein mCherry-Sec61β (control); PRG1-mCherry or the epilepsy-associated mutant PRG1-R346T-mCherry showing that PRG1 but not the mutant affects vesicle mobility.

**Movie S4: (**Related to Fig. 4d) 1 min movie (1 frame s^-1^) of DIV10 hippocampal rat neurons infected with shCT or shRNA reducing PRG1 expression (cyan) and overexpressing Syt1-mNeon (grey) showing that PRG1 silencing increases vesicle mobility at DIV10.

**Movie S5:** (Related to Fig. 6) Cryo-EM generated tomograms of tethers observed at DIV5 and DIV10 connecting the ER and the vesicle membrane. One example for each type of vesicle is shown.

**Movie S6:** (Related to Fig. 7f,g) 1 min movie (3 frames s^-1^) of DIV10 hippocampal rat neurons infected with shCT or shRNA reducing PRG1 expression (magenta Fig. 7f) and labeled with the Fluo4-AM calcium probe (green Fig. 7f and in inverted grey here) revealing that PRG1 silencing increases collective calcium bursts.

## Author contribution

G.K.V. and M.S. conceived the study and designed experiments. M.S. performed experiments. J.B.M. performed cryo-EM imaging and analysis. R.A. quantified the trafficking directionality and the fluo4-AM calcium assay. M.S. and G.K.V. wrote the manuscript. R.A. and J.B.M. reviewed the manuscript.

## Acknowledgments

We would like to acknowledge the Biochemistry Krios Electron Microscopy Facility at CU Boulder especially Dr. Erik Hartwick and Garry Morgan for their outstanding assistance. In addition, excellent support was provided by both the CU Boulder Center for Cryo Electron Tomography (CCET, Sarah Zimmermann) and the Midwest Center for Cryo-Electron Tomography (MCCET, Dr. Anna Ratliff).

## EXPERIMENTAL MODEL AND SUBJECT DETAILS

Dissociated embryonic (E18) rat hippocampi were purchased from Brain Bits (TransnetYX, Springfield, IL, SDEDHP). Cells were pelleted, counted and seeded at a density of 30,000 cells/cm^2^ on poly-L-lysine-coated (Sigma, P4707) 35 mm imaging dishes (Cellvis) for live imaging experiments or coverslips (Carolina, 633009) for fixed sample experiments. Neurons were cultured in Neurobasal-plus medium (GIBCO, A3582901) supplemented with 1xB27-plus (GIBCO, A3582801) and 1x GlutaMax (GIBCO, 35050-061). Hippocampal neurons were grown for the indicated days in vitro (DIV) before use. COS-7 cells (ATCC, CRL-1651) were cultured in DMEM (GIBCO, 12430-054) supplemented with 10% FBS and 1% Pen-Strep. HEK 293T cells (ATCC, CRL-3216) were cultured in DMEM (GIBCO, 12430-054) supplemented with 10% FBS and 1 mM sodium pyruvate. All cultures were maintained at 37°C and 5% CO2 in a humidified incubator.

## METHOD DETAILS

### Plasmids and reagents

JF646 Halo-tag ligand (Lavis lab, Janelia Research Campus;(Grimm et al., 2017)) was resuspended, stored, and used as indicated by the vendor. This study used the following published plasmids: mNeonGreen-Sec61β, mCherry-Sec61β, BFP-KDEL, mCherry-KDEL, mCherry-Rab5, pLV-mNeon-TurboID, RA-Sec61β, GB-C1,GB-Drp1, NpY-mNeon-Green, pLV-Kif5a-myc, pLV-Kif5b-myc, pLV-Kif5c-myc, GFP-Protrudin, BFP-Rab7a, BFP-Rab5, EGFP-Rab5 and EGFP-Rab7a. The following plasmids were purchased on Addgene : pEF1α-Syt1-Halo, pCl-pHluorin-Syt4 and pMDLg-RRE, pCMV-VSV-G and pRSV-REV for virus production. The following backbones available on Addgene were also used for cloning: Halo-N1, mNeonGreen-C1, mNeonGreen-N1, mCherry-C1, PlKO.1. Rat Syt1 was PCR amplified from the Syt1-Halo Addgene plasmid and cloned into the pLV-mNeon-TurboID backbone using XbaI/BamH1 sites (Fw primer: CTAGTCTAGACCACCATGGTGAGTGCCAGTCA; Rv primer: GATCGGATCCGCCTTCTTGACAGCCAGC) to obtain the pLV-Syt1-mNeon-TurboID construct. to obtain Syt1-GB, Syt1 was amplified with the restriction sites NheI/XhoI (Fw primer: CTAGGCTAGCCCACCATGGTGAGTGCCAGTCA; Rv primer: TCGACTCGAGCGGATCCCTTCTTGACA), GB was amplified with XhoI/BamHI restriction sites (Fw primer: TCGACTCGAGCCACCATGGCCACCATCAAAGA; Rv primer: GATCGGATCCTCACTTGTACCGCTCGTCCA) and both fragments were cloned into the GB-Drp1 vector digested NheI/BamHI, in replacement of GB and Drp1. The same way, Rat Syt4 was PCR amplified from the Addgene pCl-pHluorin-Syt4 plasmid and cloned into the digested XbaI/AgeI pLV-mNeon-TurboID backbone using NheI/Age1 sites (Fw primer: CTAGGCTAGCCCACCATGGCTCCTATCACCACCAG; Rv primer: TCGAACCGGTGCACCATCACAGAGCATATG) XbaI and NheI being compatible. To obtain Syt4-GB, Syt4 was PCR amplified with the restriction sites NheI/SalI (Fw primer: CTAGGCTAGCCCACCATGGCTCCTATCACCACCAG; Rv primer: TCGAGTCGACACCATCACAGAGCATATG), GB was amplified SalI/BamHI (Fw primer: TCGAGTCGACCCACCGATGGCCACCATCAAAGA; Rv primer: GATCGGATCCTCACTTGTACCGCTCGTCCA) and both fragments were cloned into the GB-Drp1 vector digested NheI/BamHI, in replacement of GB and Drp1.To obtain the Syt4-Halo plasmid, Syt4 was PCR amplified from the Addgene pCl-pHluorin-Syt4 plasmids and cloned into a Halo-N1 backbone using the NheI/AgeI sites (Fw primer: CTAGGCTAGCCCACCATGGCTCCTATCACCACCAG; Rv primer: TCGAACCGGTGCACCATCACAGAGCATATG). Syt1 and Syt4 were first PCR amplified and cloned in a mNeon green-N1 backbone using respectively NheI/BamHI (Fw primer: CTAGGCTAGCCCACCATGGTGAGTGCCAGTCA; Rv primer: GATCGGATCCGCCTTCTTGACAGCCAGC) and NheI/AgeI restriction sites (Fw primer: CTAGGCTAGCCCACCATGGCTCCTATCACCACCAG; Rv primer: TCGAACCGGTGCACCATCACAGAGCATATG). Syt1-mNeon, Syt4-mNeon and NpY-mNeon were PCR amplified and cloned into the the pLV-mNeon-TurboID in replacement of mNeon-TurboID using XbaI/BclI restriction sites (fw primer Syt1-mNeon: TCGATCTAGACCACCATGGTGAGTGCCAGTCA; Fw primer Syt4-mNeon: TCGAGCTAGCCCACCATGGCTCCTATCACCAC; Fw primer NpY-mNeon: TCGATCTAGACCACCATGCTAGGTAACAAGCG; Rv primer: TCGATGATCATTACTTGTACAGCTCGT) to obtain pLV-Syt1-mNeon, pLV-Syt4-mNeon and pLV-NpY-mNeon. EF1a-mNeon-KDEL was obtain by PCR amplification of mNeon-KDEL cloned into the pEF1a-Syt1-Halo using the EcoRI/XmaI sites (fw primer: AATTGAATTCCCACCATGAAGCTCTCCCTGGT; Rv primer: CCGGCCCGGGTTATCATAGCTCGTCTTTCTTGTACAGCTCGTCCA) in replacement of Syt1-Halo. The plasmid Ef1α-PRG1-mCherry was ordered custom and codon optimized by Twist Bioscience and a 450 base pair sequence of the natural promoter of PRG1 was added between the Ef1α promoter and the coding PRG1 sequence by Gibson assembly (NEB : E5510S) This addition improved the expression but did not change PRG1 ER localization (Geist et al., 2012). IDT DNA provided the double strand DNA fragment : (gBlocks™ Gene Fragments Entry + 5’ Phosphorylation: TCGAATTCCGGCCAGCTCCCAGTGAATTCCTTCTATGGCCTACTTGTCAGTATGAAATC TGAGTTTTAATTTTGCACAGGTGGAGGTCTCTTTTGCTATGGGTAAGGTGGATAGCGGT GCCCAGCAAGCTCCTGCTCTCTAGAAAGGCAACAAGAACCCCAGGTAGGAGACCAGGT GCCTTGGCTCCTCAGCCTTGCTTCTGCAGAAACCAGGAGTGCCTCCCCCCACTATTTTT GACGTCAAGCTCAGACAACCAGCAGAGGAGCCTCACAGCTTGGGCGGTGGAGAGCCC AGGGAGAGTGGCAGGGAGGGGAAGCCATCACAGCAACAGCTTGGAGGGGGAGCTGG CTATCACTTTCCCTCAAAATACGGACTAAAACCCGGCTGAAGAAGACCTGCGGGATCAG GGAAGCGCCGGGTTACTGCAAAGAAGGGCGGGGAAAAGGAGGGGGCGCTGCATGCA AAG). All deletion or point mutants were subcloned from the full-length sequence using the Q5 Site-Directed Mutagenesis Kit (NEB, E0554). Full-length rat PRG1-mCherry was PCR amplified from the Twist plasmid and cloned into the pLV-mNeon-TurboID backbone in replacement of mNeon-TurboID using XbaI/BclI sites (Fw primer: TCGATCTAGACCACCATGCAAAGGGCTGGAAG; Rv primer: TCGATGATCATCACTTATAGAGTTCGT). Rat PRG2 was PCR amplified from rat hippocampus cDNA and cloned into the pLV-PRG1-mCherry plasmid using XbaI/AgeI sites (Fw primer: TCGATCTAGACCACCATGATTGCTAAGAAGGAGAAGA; Rv primer: TCGAACCGGTCCGTCCTGGTACCTCCTGGC) in replacement of PRG1. For Cos7 cell transfections, PRG1 was also sub-cloned into the CMV-mNeonGreen-N1 backbone in the full-length version or the ΔC-terminus version using EcoRI/AgeI sites (Fw primer: AATTGAATTCgCCACCATGCAAAGGGCTGGAAG; Rv primer full-length: CCGGACCGGTCCATCTTTGTAGGGCCGAG; Rv primer ΔC-terminus: CCGGACCGGTCCAGAGGGATCCTGGTTAA) and in the CMV-mNeonGreen-C1 backbone in both versions as well using HindIII/EcoRI (Fw primer: AGCTAAGCTTCCATGCAAAGGGCTGGAAG; Rv primer full-length: AATTGAATTCTTAATCTTTGTAGGGCCGAG; Rv primer ΔCterminus: AATTGAATTCTTAAGAGGGATCCTGGTTAA). For knock-down experiments, we generated a shRNA-PRG1-mCherry plasmid by cloning the mCherry cassette from a mCherry-C1 backbone using BamHI/KpnI restriction sites in the PlKO.1 vector (Fw primer: CGGGATCCCCACCATGGTGAGCAAGGGCGA; Rv primer: GCGGTACCTTACTTGTACAGCTCGTCCA). The same construct was used to clone the double strand RNA hairpin using the AgeI/EcoRI sites right after the U6 promoter (Control fw: CCGGCCTAAGGTTAAGTCGCCCTCGTTCAAGAGA CGAGGGCGACTTAACCTTAGGTTTTT; Rv: AATTCCTAAGGTTAAGTCGCCCTCGTCTCTTGAA CGAGGGCGACTTAACCTTAGG)(Sh PRG1 Fw: CCGGTGGGTGAAGGGATTCTGTATTCAAGAGATACAGAATCCCTTCACCCATTTTT; Rv: AATTTGGGTGAAGGGATTCTGTATCTCTTGAA TACAGAATCCCTTCACCCA). pLV-Kif5a-myc was used to generate all Kif5a mutants using the Q5 Site-Directed Mutagenesis Kit (NEB, E0554): pLV-Kif5a-1-338-myc (Fw: GAACAAAAACTTATTTCTGAAGAAG; Rv: AGTCAACTCCAAATTTACTG), Kif5a-339-1032-myc (Fw: CTGAGCAGTGGAAGAAG; Rv: CCATTCTAGAGTCGGTG), Kif5a-1-822-myc (Fw: GCGAACAAAAACTTATTTCTGAAG; Rv: TCCCCCCACTGTCTTC). pLV-Kif5a-1-822-myc was used to generate Kif5a-339-822-myc (Fw: CTGAGCAGTGGAAGAAG; Rv: CCATTCTAGAGTCGGTG). EGFP-Rab5 and EGFP-Rab7a were cloned into the pLV-mNeon-TurboID to replace mNeon-TurboID using AgeI/BclI restriction sites to obtain pLV-EGFP-Rab5 and pLV-EGFP-Rab7a.

### Transfections

Neurons were grown on 35 mm imaging dishes (Cellvis, D35-20-1.5-N) and transfected with the indicated plasmids using Lipofectamine 2000 (Thermo Fisher Scientific, 11668019), according to manufacturer instructions. Briefly, two separate 125 µL reactions were assembled in Opti-MEM (Invitrogen, 31985-088), one with 5 µL of Lipofectamine 2000 and the other with 2 µg of plasmid mixture in total. After 5 min, reactions were combined and kept at room temperature for 20 min, after which the mixture was added to cells and incubated for 24 hours. For cell lines, reactions were prepared in Opti-MEM (Invitrogen, 31985-088). Transfection was achieved using Lipofectamine 3000 (Thermo Fisher Scientific, L3000150) and plasmids were mixed with Reagent 3000 at 5 µL/mL. Cells were incubated with the transfection mixture overnight. Neurons and cell lines were imaged 24 h after transfection in their respective media.

### Knockdown with shRNA

For shRNA experiments, neurons were infected at DIV4 with lentiviruses carrying control or shRNA targeting rat PRG1. If necessary, cells were transfected as indicated in the transfection section at DIV8 with the vesicle markers and finally imaged at DIV10. If not, they were used at DIV10 for fixation or Fluo4-AM calcium imaging experiments. Samples were also collected from 6 well plates for knockdown verification by immunoblot and qPCR.

### Reverse transcription and quantitative PCR (RT-qPCR)

Total RNA was isolated using the Monarch Total RNA Miniprep Kit (New England Biolabs, T2010S) following the manufacturer’s instructions, including the on-column DNase I treatment to remove residual genomic DNA. RNA quantity and purity were assessed by spectrophotometry (NanoDrop, Thermo Fisher Scientific). For each sample, 500 ng to 1 µg of RNA was reverse-transcribed using the LunaScript RT Master Mix Kit (New England Biolabs, E3025L) in a final reaction volume of 20 µL, according to the supplier’s recommendations. qPCR was performed using the Luna Universal qPCR Master Mix (New England Biolabs, M3003X) on a 7500 qPCR fast System (Applied Biosystems). Each 20 µL reaction contained 4 µL of diluted cDNA and gene-specific primers purchased from IDT DNA: Rat ActinB: Rn.PT.39a.22214838.g, Rat Lppr4 (PRG1): Rn.PT.58.19098303, Rat Lppr3 (PRG2): Rn.PT.58.35042039 and Rat Zfyve27 (Protrudin): Rn.PT.58.44751837. All reactions were run in technical duplicate and at least 4 biological replicates. Cycling conditions consisted of an initial denaturation at 95°C for 1 min, followed by 40 cycles of 95°C for 15 s and 60°C for 30 s, during which fluorescence acquisition was performed. At the end of each run, melt-curve analysis was used to confirm amplification specificity and absence of primer-dimers. Relative gene expression levels were calculated using the ΔΔCt method, with normalization to the housekeeping gene ActinB. Primer efficiency was evaluated using cDNA dilution curves, and only primer pairs exhibiting efficiencies over 95% were included in the analysis.

### Sample preparation, immunoblot analysis, and immunocytochemistry

For immunoblot (IB) experiments, cell cultures were homogenized in RIPA-like buffer (225 mM mannitol, 75 mM sucrose, 0.1 mM EGTA, 0.1% SDS, 30 mM Tris-HCl, pH7.4) supplemented with protease inhibitor cocktail (Complete Mini EDTA-free protease inhibitor cocktail tablets, Roche,11836170001). Samples were then mixed with Laemmli sample buffer (125 mM Tris-HCl, 4% SDS, 20% glycerol, 5% 2-mercapto-ethanol, bromophenol blue, pH 7.4) boiled and resolved by SDS-Polyacrylamide Gel Electrophoresis (SDS-PAGE) on 4-20% gels (BioRad, Criterion TGX pre-cast gels, 5671094) and transferred to polyvinylidene fluoride (PVDF) membranes (Millipore, IPVH00010). PVDF membranes were blocked with 5% nonfat dried milk, in Tris-buffered saline (TBS) pH 7.4 with 0.1% Tween20 and probed with the indicated antibodies. Primary antibody binding was detected with horseradish peroxidase-conjugated secondary antibodies and visualized by chemiluminescence (Thermo FisherScientific, SuperSignal West FemtoPLUS, 34096).

For immunocytochemistry (ICC) experiments, cells were grown on pre-coated coverslips, fixed with 2% paraformaldehyde in PBS for 15 min at 37°C. Samples were then quenched at room temperature for 10 min with 50 mM ammonium chloride, washed with PBS, and permeabilized with blocking buffer (10% Horse serum and 1% BSA in PBS) containing 0.1% Triton X-100 for 10 min at room temperature. Cells were washed and incubated again for 30 min at room temperature with the blocking buffer before being incubated overnight at 4°C with primary antibodies diluted in the same blocking buffer. After three washes, cells were incubated with fluorophore-conjugated secondary antibodies for 2 h at room temperature. Cells were washed again three times with blocking buffer before mounting.

Primary antibody dilutions used: Anti-PRG1 Mouse monoclonal 1/500; Anti-VAP-B Rabbit polyclonal 1/750; Mouse-anti-Syt1 Mouse monoclonal 1/100; Anti-Syt4 Rabbit polyclonal 1/600; Anti-GAPDH Rabbit polyclonal 1/2500; Anti-β-Actin 1/1000; Rabbit monoclonal; Anti-PRG2 Rabbit polyclonal 1/500; Anti-Synaptophysin Mouse monoclonal 1/2000; Anti-Bassoon Mouse monoclonal 1/500 and Anti-Histone H3 rabbit polyclonal 1/1000 for western Blot and IF; Anti-Homer Rabbit polyclonal 1/500; Anti-V5 1/2000; Anti-Kif1a 1/750; Anti-Kif5a 1/500; Anti-myc 1/1000; Anti-βIII-tubulin 1/5000; Anti-mCherry 1/750; Anti-Protrudin 1/500. Secondaries for immunoblots: Goat anti-Mouse IgG or Goat anti-Rabbit IgG 1/5000, Anti-streptavidin-HRP 1/6000; Goat anti-Rabbit Highly Cross-Absorbed 1/2500; Goat Anti-Rat-AF647 1/2500 and Donkey Anti-Rabbit-AF647 1/2500. Secondaries for immunocytochemistry: Anti-streptavidin-AF555 1/5000; Anti-mCherry-AF594 1/400; Donkey-Anti-Mouse-AF647 and Donkey-Anti-Rabbit-AF488 1/800.

### Co-immunoprecipitation

DIV10 primary rat neurons were washed two times with ice-cold PBS and lysed in buffer containing 25 mM Tris-HCl, pH 7.4, 150 mM NaCl, 5% glycerol, 0.5% NP-40 and 1mM EDTA supplemented with a protease inhibitor cocktail (Complete Mini EDTA-free protease inhibitor cocktail tablets, Roche, 11836170001) and Phophatase inhibitor cocktail (PhosSTOP, Sigma, 4906845001). Lysates were nutated for 30 min at 4°C and then cleared by 1000g centrifugation 5 min at 4°C. An aliquot of each cleared lysate was retained as input. The remaining lysate was incubated with mouse anti-PRG1 antibody (R&D Systems, MAB2874) or an equivalent amount of mouse IgG isotype control (R&D Systems, MAB002) overnight at 4°C with continuous rotation. Protein A/G beads (Thermofisher, 88802) were then added, and samples were incubated for an additional hour at 4°C. Beads were collected and washed three times with ice-cold lysis buffer. Bound proteins were eluted by adding 2x Laemmli sample buffer and heated at 95°C for 5 min. Inputs and immunoprecipitated samples were resolved by SDS–PAGE and analyzed by immunoblotting or mass spectrometry.

HEK293T cells were transfected with designated combinations of plasmids and grown for 24 hours. Cultures were pelleted in cold 1x PBS. Cells were resuspended in IP lysis buffer (25 mM Tris-HCl pH 7.4, 150 mM NaCl, 1 mM EDTA, 0.5% NP-40 and 5% glycerol) supplemented with a protease inhibitor cocktail (Complete Mini EDTA-free protease inhibitor cocktail tablets, Roche, 11836170001) and Phophatase inhibitor cocktail (PhosSTOP, Sigma, 4906845001). Lysates were nutated for 30 min at 4°C and then cleared by high-speed centrifugation for 10 min at 4°C. The supernatant was incubated 1 hour at 4°C with mCherry-Trap agarose magnetic beads (Chromotek, rtma). Beads were pelleted, washed 3 times with ice-cold lysis buffer, resuspended in 2x Laemmli buffer and heated at 95°C for 5 min. Samples were resolved on an SDS-PAGE gel and blotted for specified antibodies.

### TurboID proximity biotinylation

Rat neurons were transfected with the designated TurboID-fused constructs at DIV6. At DIV7, cultures were treated with 500 µM biotin (Sigma, B4501) for 15 minutes, washed 3 times with Neurobasal maintenance media and once with 1x PBS, and were scraped with RIPA-like lysis buffer (225 mM mannitol, 75 mM sucrose, 0.1 mM EGTA, 0.1% SDS, 30 mM Tris-HCl, pH 7.4) supplemented with protease inhibitor cocktail (Complete Mini EDTA-free protease inhibitor cocktail tablets, Roche,11836170001). Cell lysates were centrifuged at max speed for 10 min to remove unsolubilized material, and the supernatant was vortexed. All samples were flash frozen in liquid nitrogen and stored at -80°C until analyzed by mass spectrometry at the Sanford Burnham Prebys Medical Discovery Institute proteomics facility.

### SoRa live-cell microscopy

Cells were imaged with a Yokogawa CSU-W1 SoRa spinning disk confocal, which is built on a Ti2 inverted microscope and contains a Hamamatsu ORCA-FusionBT sCMOS camera (Nikon). In all cases, a 60x (1.42 NA) Plan Apo objective was used with 2.8x SoRa magnification giving a pixel size of 40 nm. The fluorescence channels were 488 nm excitation (FITC emission filter, BP 525/36), 561 nm excitation (TRITC emission filter, BP 605/52), 647 nm excitation (Cy5 emission filter, BP705/72) and 405 nm excitation (BFP emission filter, BP 455/50). All timelapse images were deconvolved with a maximum of 20 iterations of a Richardson-Lucy algorithm in NIS-Elements software (Nikon). For live-cell experiments, dishes were kept at 37°C in a humidified atmosphere with 5% CO2.

### Large image acquisition and stitching

Neuronal cultures prepared for cryo-electron microscopy were imaged on a Nikon Ti2 inverted microscope equipped with a Yokogawa CSU-X1 spinning disk confocal scanner and a 100x/1.45 NA Plan Apo λ DIC N2 oil objective. An additional ∼1.42x intermediate magnification yielded a calibrated pixel size of 0.113 µm. mCherry was excited with a 561 nm laser (OBIS, Coherent) and images were recorded with an Andor iXon Ultra 897 EMCCD camera. Large-field images were acquired in Micro-Manager (version 2.0) as 5 × 5 grids of tiles with ∼10% overlap, with the stage moving between tile positions. Grid start positions were selected by eye using DIC, blinded to mCherry fluorescence. Three separate grids were acquired per condition.

### Software and code availability

Tiles were stitched with a custom Python script (NumPy 1.26.4, tifffile 2024.6.18, matplotlib 3.9.2). Code development was assisted by Claude (Sonnet 5; Anthropic, San Francisco, CA), an AI coding assistant.

### Cryo-EM sample preparation and imaging

Quantifoil 2/2, 2/1 or 1.2/1.3 gold 200 mesh grids (EM Sciences) were glow discharged, UV sterilized and placed into 35mm imaging dishes. Grids were coated overnight with poly-L-lysine at 4°C. The following day, dishes were rinsed thoroughly with Neurobasal media, placed in Neurobasal supplemented media, and cells were seeded at low density (20,000 cells/ cm^2^). At DIV5 or 10, 10nm gold particles (fiducials) suspended in Neurobasal media were applied to the cell side of the grid, and grids were then blotted from the non-cell side for 5-7 seconds and plunge frozen in liquid ethane using a Leica EM GP2 plunge freezer. Tilt series of intact axons were acquired with SerialEM, Tomo or Tomo5 on a Titan Krios G3i, G4 or 5 equipped with a Falcon 4i direct electron detector (DED) and a Selectris or Selectris X energy filter operating at a 10eV slot width (Thermo Fisher). Tilt series were also acquired with SerialEM using a Glacios with a Gatan K3 DED (Thermo Fisher). Images were collected at 2 or 3 degree increments from +/-56-60 degrees using a dose symmetric tilt scheme for a total dose of 97-117 e^-^/Å^2^. Defocus values ranged from -4 to -7, and the acquisition magnification was 33000x with a pixel size of 3.726 or 3.83 Å/px (Glacios magnification acquisition was 13500x with a pixel size of 2.922 Å/px).

### Lentiviral particle production

HEK293T cells were seeded on 10 cm dishes and grown to 80% confluence. Cultures were co-transfected with pVSV-G, pREV, and pRRE helper plasmids plus a designated shuttle vector carrying the sequence of interest. Lentivirus-containing media was collected at 48 and 72h, pooled, and filtered. The supernatant was aliquoted in 1.5 mL eppendorf tubes and spun down at max speed (16000 g) on a table-top centrifuge for 2 h at 4°C to pellet lentiviral particles. The supernatant was discarded and dry lentiviral pellets were stored at -80°C until use.

### Fluo4-AM calcium assay

Rat hippocampal neurons were plated in a dark, glass bottom, pre-coated 96 well plate (25,000 cells per well). Cells were infected at DIV4 with shRNA containing viruses and used at DIV10. The Neurobasal maintenance media was refreshed with 5 µM of the Fluo4-AM probe (APEXBio, B8807) for 30 minutes at 37°C. Cells were washed once with Neurobasal maintenance media and once with artificial cerebrospinal fluid (ACSF, Tocris, 3525). Cells were incubated again for 30 min at 37°C in ACSF before being imaged. This experiment was performed using the 20x PLAN APO λD NA = 0.80 air objective of the SoRa microscope without intermediate magnification and the Hamamatsu Orca BT Fusion sCMOS camera yielding a pixel size of 0.33 µm per pixel. This microscope was used to record 1 min movies at 3 images s^-1^ for the green channel (calcium Fluo4-AM probe). Another image was acquired after the movie to image both the green and the red channel in order to detect shRNA-mCherry positive cells. The experiment was done 3 times. For each experiment, 4 movies from 4 different wells were recorded per condition. Emission filters are described above.

## QUANTIFICATION AND STATISTICAL ANALYSIS

### Image processing and quantifications

Fluorescence microscopy images were processed using Fiji (Schindelin et al., 2012). All line scans were generated with the ‘‘Plot profile’’ tool of FIJI. Kymographs were generated using the ‘‘Multi Kymograph’’ tool.

### ER-vesicle overlap

Syt1- or Syt4-positive vesicles were segmented by thresholding to generate binary masks. These masks were combined with the ER channel (mNeon-KDEL) to measure ER fluorescence intensity at individual vesicle locations. Vesicles displaying a mean ER fluorescence intensity above the background threshold were considered ER-overlapped. As a control for random colocalization, the vesicle mask was horizontally flipped and the analysis was repeated. The percentage of ER-associated vesicles was calculated as the number of vesicles above the background threshold divided by the total number of analyzed vesicles. For each cell, 3 areas of 6x6 µm were picked along the axon and analyzed.

### ddFP

ddFP fluorescence was measured in 3 × 3-pixel ROIs positioned at ER–vesicle overlaps and in ER-only regions. Local background was subtracted, and corrected fluorescence values were normalized to the mean intensity of all ROIs within the same cell. An ER–vesicle overlap was classified as ddFP-positive when its normalized intensity exceeded the maximum value measured in ER-only regions from the same cell.

### Vesicle trafficking

SVP and DCV trafficking were scored from Trackmate (LoG detector-Advanced Kalman Tracker) for each movie (Ershov et al., 2022). The quality threshold was manually optimized for each cell because of over-expression level variability. The total distance travelled by each vesicle was plotted. Tracks shorter than 2 frames were discarded. Vesicles were classified as ‘‘Mobile’’ (displacement ≥ 1.5 µm) or ‘‘Immobile’’ (displacement < 1.5 µm), and the numbers for both categories were normalized to the total number of vesicles analyzed. All vesicle-trafficking analyses were restricted to the first 45 s of each recording to ensure an identical observation period across experiments. Longer recordings were truncated to 45 s before tracking. In DIV10 experiments, the mean speed and maximum instantaneous speed were extracted for each track with a displacement exceeding 1.5 µm. Mean track speed reflects the effective speed over the entire trajectory and is therefore influenced by pauses, whereas maximum instantaneous speed estimates the velocity reached during active movement. Values were then averaged at the cell level for statistical analysis.

### Directionality analysis

For directionality, kymographs were generated and directionally separated using the KymoClear 2.0 plugin in Fiji (Mangeol et al., 2016). Only axons for which the orientation could be unambiguously determined were included. Anterograde, retrograde, and stationary signals were segmented independently using ilastik. Because stationary signals can be retained in the directionally filtered anterograde and retrograde images, the stationary segmentation mask was subtracted from both directional masks to prevent the same signal from being counted in multiple categories. The number of pixels assigned to each category was normalized to the total number of segmented pixels within the corresponding kymograph.

### Quantification of protein expression

To quantify the relative fluorescence expression of the different PRG1 constructs, neurons were segmented in Fiji following background subtraction and Gaussian blur filtering. The resulting masks were applied to the corresponding unprocessed images to measure the mean fluorescence intensity within each neuron. Mean background fluorescence was measured from a cell-free region of the same image. To account for differences in illumination settings, laser output was calibrated by measuring the power at the objective back focal plane with a Thorlabs power meter at four laser-power percentages spanning the range used for image acquisition. The displayed laser-power percentages were plotted against the measured output in mW, and a linear regression was performed in Python using the linregress function from the SciPy package. The resulting regression equation was used to estimate the laser power delivered for each acquired image. Relative expression was calculated by dividing the background-corrected mean neuronal fluorescence by the calibrated laser power, expressed in mW, and the exposure time.

### Perinuclear Rab7a localization

Three-channel Z-stacks of COS-7 cells were analyzed using custom Fiji macros. Cell and nuclear outlines were manually delineated, and an extracellular region was selected to estimate background fluorescence.

Rab7a-positive structures were segmented independently on each Z-plane. A Gaussian-smoothed image (σ = 15 pixels) was subtracted from a lightly smoothed image (σ = 1 pixel) to suppress diffuse fluorescence. Pixels were retained when the resulting local contrast exceeded the extracellular background mean plus six standard deviations and reached at least 30% of the local background-corrected Rab7a signal. Connected regions smaller than 40 pixels were removed.

The cytoplasmic region was defined as the cell ROI excluding the nuclear ROI. A two-dimensional Euclidean distance map was calculated from the nuclear mask, and the perinuclear region comprised cytoplasmic pixels located within 8 µm of the nuclear boundary. This region was applied to every Z-plane.

For each cell, perinuclear Rab7a localization was calculated as: Perinuclear Rab7a (%) = 100 × (Rab7a-positive pixels within the perinuclear region) / (Rab7a-positive pixels within the entire cytoplasm).

Pixels outside the cell or within the nuclear ROI were excluded. All retained Rab7a-positive pixels contributed equally. Measurements were combined across the complete Z-stack to obtain one percentage per cell. Cell-level percentages were averaged within each independent experiment before statistical comparison.

### Quantification of putative synapses

Immunofluorescence experiments were analyzed with FIJI, 50 µm-long axonal sections were analyzed. In these segments, “synapse” events were manually counted and measured. Puncta were classified as putative synapses when Homer and Bassoon signals overlapped by at least one pixel and the size of the pattern was ≥ 0.2 µm. Events located more than 2 µm away from the mCherry signal (corresponding to shRNA-expressing neurons) were excluded from the analysis.

## Fluo4-AM analysis

For Fluo4-AM calcium experiment quantifications, images were preprocessed in two steps by cropping the central 1864x1864 pixels and performing flat-field correction. A CellProfiler pipeline was used to segment neuron cell bodies based on mCherry signal (Stirling et al., 2021). First, images were smoothed using a Gaussian filter with an 8-pixel diameter. Cell bodies were then detected using the Cellpose ‘cyto’ pre-trained model. Resulting objects were shrunk by one pixel, to ensure separation of objects. Next, Fluo4-AM mean intensity was measured in each object over the entire timelapse. The measurements for each object (or cell) were analyzed using a custom python script. The SciPy python library (specifically the ‘find_peaks’ algorithm) was used to find peaks with a prominence of either 0.1 or 0.01 in the timelapse images (Virtanen et al., 2020). Peaks were counted, graphed, and statistics calculated using Graphpad Prism.

### Cryo-ET data analysis

The IMOD software package was used to generate tomograms from tilt series of intact axons using gold fiducials for alignment. Tomograms were binned by 2 and Nonlinear Anisotropic Diffusion (NAD) filtering was applied to increase contrast for segmentation (Kremer et al., 1996; Mastronarde & Held, 2017). No additional denoising procedure was applied. To determine the distance between SVs or DCVs and the ER, the shortest membrane-to-membrane distance was measured by drawing contours between the organelles in the slicer feature of IMOD. Vesicle diameter measurements were obtained by drawing contours from the outer edges of the SV or DCV membranes. Visible tether-like densities were scored when an electron-dense structure visibly bridged the ER and vesicle membranes, and tether length was measured between the two membranes along the visible density, following irregular shapes. Each vesicle less than 30 nm away from the ER was extracted for detailed analyses and its distance to the ER as well as its proximity to a microtubule were reported on the graphs in Figure 6. The shortest distance between the vesicle membrane and the surface of the nearest microtubule was measured, and vesicles located within 30 nm of both the ER and a microtubule were classified as tripartite ER-vesicle-microtubule junctions.

### Infection efficiency analysis

To determine the infection efficiency of the mCherry-expressing shRNA constructs used for cryo-EM experiments, corresponding DIC and mCherry images of DIV10 neuronal cultures infected with either shCT or shPRG1 were acquired and stitched to generate large fields of view. Neuronal cell bodies were identified and counted from the DIC images independently of the mCherry channel. Infection efficiency was calculated for each stitched field as the percentage of DIC-identified neurons exhibiting somatic mCherry fluorescence at least twofold above the background signal. All the knockdown experiments were also verified by WB and RT-qPCR.

### Statistical analysis

In all superplots illustrating vesicle trafficking quantification, each point represents one track analyzed, and each experiment was performed at least 3 times independently. Every mean of each experiment is shown as a colored triangle with n=1 Orange; n=2 Blue and n=3 Pink etc. Statistical analyses were performed on GraphPad Prism 10 using the means of independent experiments. Student’s paired or unpaired t-tests were used, as indicated. When testing significance for more than two conditions, one-way ANOVA was performed. Details of statistical analysis and the number of cells quantified in each experiment are indicated in the figure legends.

## Data availability

All data supporting the findings of this study are available within the article, its Supplementary Information, and the accompanying Source Data files. Source Data include the numerical data underlying all graphs and statistical analyses, unprocessed immunoblots, and mass spectrometry datasets. The microscopy movies are provided as Supplementary Movies.

| **REAGENT or RESOURCE** | **SOURCE** | **IDENTIFIER** |
|---|---|---|
| **Antibodies** |  |  |
| Anti-PRG1 Mouse monoclonal | R&D systems | Catalog#mab2874 |
| Anti-VAP-B Rabbit polyclonal | Sigma | Catalog#HPA013144 |
| Anti-Syt1 Mouse monoclonal | DSHB | Catalog#mAb48(asv48) |
| Anti-Syt4 Rabbit polyclonal | Genetex | Catalog#GTX32906 |
| Anti-GAPDH Rabbit polyclonal | Sigma | Catalog#G9545 |
| Anti-β-Actin Rabbit monoclonal | Genetex | Catalog#GTX637675 |
| Anti-PRG2 Rabbit polyclonal | Proteintech | Catalog#25936-1-AP |
| Anti-Synaptophysin Mouse monoclonal | Synaptic Systems | Catalog#101011 |
| Anti-Histone H3 rabbit polyclonal | Abcam | Catalog#Ab1791 |
| Anti-Bassoon Mouse monoclonal | Neuromab | Catalog#75-491 |
| Anti-Homer Rabbit polyclonal | Synaptic Systems | Catalog#160003 |
| Anti-V5 | Thermofisher | Catalog#R960CUS |
| Anti-Mouse IgG Isotype Control | R&D systems | Catalog#mab002 |
| Anti-Kif1a Mouse monoclonal | BD Transduction Laboratories | Catalog#612094 |
| Anti-Kif5a Rabbit polyclonal | Abcam | Catalog#Ab5628 |
| Anti-mCherry Rat 16d7 monoclonal | Invitrogen | Catalog#740003T20UG |
| Anti-βIII-tubulin Mouse monoclonal (TuJ1) | R&D systems | Catalog#MAB1195 |
| Anti-Myc-Tag (9B11) Mouse monoclonal | Cell signaling | Catalog#2276S |
| Anti-Protrudin Rabbit polyclonal | Proteintech | Catalog#12680-1-AP |
| Goat anti-Mouse IgG (whole molecule)-<br>Peroxidase antibody | Sigma-Aldrich | Catalog#A4416 |
| Goat anti-Rabbit IgG (whole molecule)-<br>Peroxidase antibody | Sigma-Aldrich | Catalog#A6154 |
| Goat anti-Rabbit IgG (whole molecule)- Highly cross-Adsorbed Peroxidase antibody | Invitrogen | Catalog#A16110 |
| Goat anti-Rat IgG (H+L) Highly Cross-Adsorbed Secondary Antibody, Alexa Fluor 647 (immunoblots) | Thermofisher | Catalog#A21247 |
| Donkey anti-Rabbit IgG (H+L) Highly Cross-Adsorbed Secondary Antibody, Alexa Fluor 647 (immunoblots) | Thermofisher | Catalog#A31573 |
| Anti-streptavidin-HRP | Invitrogen | Catalog#434323 |
| Anti-Streptavidin, Alexa Fluor 555 conjugate, 0.5 ml | Thermofisher | Catalog#S32355 |
| Anti-mCherry Monoclonal Antibody (16D7), Alexa Fluor 594 | Thermofisher | Catalog#M11240 |
| Donkey anti-Mouse IgG (H+L) Highly Cross-Adsorbed Secondary Antibody, Alexa Fluor Plus 647 | Thermofisher | Catalog#A32787 |
| Donkey anti-Rabbit IgG (H+L) Highly Cross-Adsorbed Secondary Antibody, Alexa Fluor 488 | Thermofisher | Catalog#A21206 |
| **Chemicals** |  |  |
| Fluo4-AM | ApexBio | Catalog#B8807 |
| Janelia Fluor 646 HaloTag Ligand | Janelia Fluor | N/A |
| Lipofectamine 3000, transfection reagent | ThermoFisher | Catalog#L3000015 |
| Lipofectamine 2000, transfection reagent | ThermoFisher | Catalog#11668030 |
| Poly-L-Lysine solution | Sigma | Catalog#P4707 |
| **Commercial assays** |  |  |
| LunaScript RT Master Mix Kit | NEB | Catalog#E3025L |
| Luna Universal qPCR Master Mix | NEB | Catalog#M3003X |
| QIAquick Gel Extraction Kit | QIAGEN | Catalog# 28706 |
| ZymoPURE Plasmid Miniprep Kit | Zymo research | Catalog#D4211 |
| SuperSignal West Femto PLUS Chemiluminescent Substrate | Thermofisher | Catalog#34096 |
| Monarch Total RNA Miniprep Kit | NEB | Catalog#T2010S |
| Q5 Site-Directed Mutagenesis Kit | NEB | Catalog#E0554 |
| **Experimental models: cell lines** |  |  |
| Cos7 | ATCC | Cat# CRL-1651, RRID:CVCL_0224 |
| HEK293T | ATCC | Cat# CRL-3216, RRID:CVCL_0063 |
| **Experimental models: Organisms/strains** |  |  |
| E18 Rat Dissociated Hippocampus | Brain Bits, Transnet YX https://tissue.transnetyx.com/ | SKU : SDEDHP |
| **Recombinant DNA** |  |  |
| EF1α-mNeonGreen-KDEL | This study | N/A |
| mCherry-KDEL | (Zurek et al., 2011) | N/A |
| BFP-KDEL | (Friedman et al., 2010) | N/A |
| mNeonGreen-Sec61β | (Wu & Voeltz, 2021) | N/A |
| mCherry-Sec61β | (Zurek et al., 2011) | N/A |
| pEF1α-Syt1-Halo | Addgene | Plasmid#197281 |
| pLV-Syt1-mNeonGreen | This paper | N/A |
| pCI-pHluorin-Syt4 | Addgene | Plasmid#184505 |
| Syt4-Halo | This paper | N/A |
| pLV-Syt4-mNeonGreen | This paper | N/A |
| NpY-mNeonGreen | (Zamponi et al., 2022) | N/A |
| pLV-NpY-mNeonGreen | This paper | N/A |
| GB-C1 | (Lee et al., 2020) | N/A |
| Syt1-GB | This paper | N/A |
| Syt4-GB | This paper | N/A |
| pCI-pHluorin-Syt4 | Addgene | Plasmid#184505 |
| GB-Drp1 | From the lab | N/A |
| RA-sec61β | (Lee et al., 2020) | N/A |
| EF1α-PRG1-mCherry | Twist | N/A |
| PRG1-mNeonGreen | This paper | N/A |
| PRG1-ΔCter-mNeonGreen | This paper | N/A |
| mNeonGreen-PRG1 | This paper | N/A |
| mNeonGreen-PRG1ΔCter | This paper | N/A |
| EF1α-PRG1-mCherry | This paper | N/A |
| EF1α-PRG1-R346T-mCherry | This paper | N/A |
| EF1α-PRG1-L348D-mCherry | This paper | N/A |
| EF1α-PRG1-R346K-mCherry | This paper | N/A |
| EF1α-PRG1-ΔCter-mCherry | This paper | N/A |
| pLV-PRG1-mCherry | This paper | N/A |
| pLV-PRG2-mCherry | This paper | N/A |
| pLV-shRNA-CT-mCherry | This paper | N/A |
| pLV-shRNA-PRG1-mCherry | This paper | N/A |
| pMDLg-RRE | Addgene | Plasmid#12251 |
| pCMV-VSV-G | Addgene | Plasmid#8454 |
| pRSV-REV | Addgene | Plasmid#12253 |
| pLV-mNeon-TurboID | (Zamponi et al., 2022) | N/A |
| pLV-Syt1-mNeon-TurboID | This paper | N/A |
| pLV-Syt4-mNeon-TurboID | This paper | N/A |
| pLV-EGFP-Rab5 | This paper | N/A |
| pLV-EGFP-Rab7a | This paper | N/A |
| AcGFP1-Protrudin (isoC) | (Hoyer et al., 2018) | N/A |
| BFP-Rab7a | (Friedman et al., 2013) | N/A |
| BFP-Rab5 | (Friedman et al., 2013) | N/A |
| pLV-Kif5a-myc | (Zamponi et al., 2022) | N/A |
| pLV-Kif5b-myc | (Zamponi et al., 2022) | N/A |
| pLV-Kif5b-myc | (Zamponi et al., 2022) | N/A |
| pLV-Kif5a-1-338-myc | This paper | N/A |
| pLV-Kif5a-339-1032-myc | This paper | N/A |
| pLV-Kif5a-1-822-myc | This paper | N/A |
| pLV-Kif5a-339-822-myc | This paper | N/A |
| **Software and algorithms** |  |  |
| Fiji | (Schindelin et al., 2012) | N/A |
| Expression atlas – EMBL-EBI | www.ebi.ac.uk | N/A |
| Trackmate | (Ershov et al., 2022) | N/A |
| Kymoclear 2.0 | (Mangeol et al., 2016) | N/A |
| CellProfiler | (Stirling et al., 2021) | N/A |
| SciPy | (Virtanen et al., 2020) | N/A |
| Heliquest | (Gautier et al., 2008) | N/A |
| DISOPRED3 | https://bioinf.cs.ucl.ac.uk/psipred/ | N/A |
| Alphafold | https://alphafoldserver.com | N/A |
| DeepTMHMM | https://services.healthtech.dtu.dk/services/DeepTMHMM-1.0/ | N/A |
| IMOD | (Kremer et al., 1996; Mastronarde & Held, 2017) | N/A |
| Graphpad Prism 10 | Version 10.5.0(673) | N/A |
| Snapgene | Version 8.2.1 | N/A |
| Adobe illustrator | Version 29.8.5 | N/A |
| Ilastik | (Berg et al., 2019) | N/A |

